# Metabolic interactions control the spread of plasmid-encoded functional novelty during microbial range expansion

**DOI:** 10.1101/2022.06.07.495077

**Authors:** Yinyin Ma, Anton Kan, David R. Johnson

**Affiliations:** Department of Environmental Microbiology, Swiss Federal Institute of Aquatic Science and Technology (Eawag), 8600 Dübendorf, Switzerland; Department of Environmental Systems Science, Swiss Federal Institute of Technology (ETH), 8092 Zürich, Switzerland; Department of Materials, Swiss Federal Institute of Technology (ETH), 8093 Zürich, Switzerland

**Author notes:** Corresponding authors: Yinyin Ma, Eawag, Department of Environmental Microbiology, Überlandstrasse 133, 8600 Dübendorf, Switzerland., David R. Johnson, Eawag, Department of Environmental Microbiology, Überlandstrasse 133, 8600 Dübendorf, Switzerland.

**Keywords:** Antibiotic resistance, Horizontal gene transfer, Conjugation, Microbial Interactions, Range expansion

## Abstract

Surface-associated microbial communities are omnipresent on Earth. As individuals grow and divide within these communities, they undergo range expansion during which different cell-types arrange themselves across space to form spatial patterns (referred to as spatial self-organization). Metabolic interactions are important determinants of the spatial self-organization process, where they direct the spatial positionings of different cell-types. We hypothesized here a previously unexplored consequence of metabolic interactions; by directing the spatial positionings of different cell-types, they also control the horizontal spread of functional novelty during range expansion. We focused on a form of functional novelty of critical importance to human health – the conjugative transfer and proliferation of plasmid-encoded antibiotic resistance. We performed range expansion experiments and spatially-explicit individual-based computational simulations with pairs of strains of the bacterium *Pseudomonas stutzeri*, where one strain was a plasmid donor and the other a potential recipient. We then imposed a competitive or resource cross-feeding interaction between them. We found that interactions that increase the spatial intermixing of strains also increase plasmid conjugation. We further directly linked these effects to spatial intermixing itself. We finally showed that the ability of plasmid recipients to proliferate is determined by their spatial positionings. Our results demonstrate that metabolic interactions are indeed important determinants of the horizontal spread of functional novelty during microbial range expansion, and that the spatial positionings of different cell-types need to be considered when predicting the proliferation and fate of plasmid-encoded traits.

## Introduction

Surface-associated microbial communities are ubiquitous across our planet and have critical roles in human health and disease, biogeochemical cycling, and biotechnology (Hall-Stoodley, Costerton et al. 2004, Singh, Paul et al. 2006, Bruellhoff, Fiedler et al. 2010, Nobile and Johnson 2015, Battin, Besemer et al. 2016). As individuals within these communities grow and divide, they exert physical forces on neighboring cells that cause communities as a whole to expand across space (referred to as range expansion) (Hallatschek, Hersen et al. 2007, Weinstein, Lavrentovich et al. 2017, Giometto, Nelson et al. 2018). During the range expansion process, different cell-types arrange themselves non-randomly across space (referred to as spatial self-organization [SSO]) (Ben-Jacob, Cohen et al. 2000, Tolker-Nielsen and Molin 2000, Smith, Davit et al. 2017). The patterns of SSO that develop depend on local environmental conditions (Gralka, Stiewe et al. 2016, Mitri, Clarke et al. 2016, Sharma and Wood 2021, Ciccarese, Micali et al. 2022), the phenotypes expressed by individuals (Rudge, Federici et al. 2013, Gralka, Stiewe et al. 2016, Smith, Davit et al. 2017, Xiong, Cao et al. 2020), cell-cell interactions (Momeni, Waite et al. 2013, Blanchard and Lu 2015, Nadell, Drescher et al. 2016, Goldschmidt, Regoes et al. 2017, Tecon and Or 2017, Kan, Del Valle et al. 2018), and cell-surface interactions (Atis, Weinstein et al. 2019, Ciccarese, Zuidema et al. 2020, Fei, Mao et al. 2020). Importantly, SSO can be an important determinant of the collective traits of communities and the evolutionary processes acting on those communities (Gralka, Stiewe et al. 2016, Nadell, Drescher et al. 2016, Weinstein, Lavrentovich et al. 2017, Giometto, Nelson et al. 2018, Kayser, Schreck et al. 2018, Bosshard, Peischl et al. 2019, Goldschmidt, Caduff et al. 2021).

Metabolic interactions between different cell-types are pervasive within microbial communities and can direct the spatial positionings of cell-types during the SSO process (Nadell, Foster et al. 2010, Momeni, Waite et al. 2013, Müller, Neugeboren et al. 2014, Nadell, Drescher et al. 2016, Goldschmidt, Regoes et al. 2017, Rodríguez Amor and Dal Bello 2019). Resource competition generally results in lower spatial intermixing of cell-types as a consequence of small effective populations at the expansion frontier that are susceptible to ecological drift (Hallatschek, Hersen et al. 2007, Excoffier, Foll et al. 2009, Hallatschek and Nelson 2010). Conversely, resource cross-feeding, where one cell-type produces a resource that can be used by others, generally results in higher spatial intermixing (Momeni, Brileya et al. 2013, Müller, Neugeboren et al. 2014, Goldschmidt, Regoes et al. 2017). This can be caused by *i*) metabolic dependencies between cell-types that counteract the effects of ecological drift at the expansion frontier (Momeni, Brileya et al. 2013, Müller, Neugeboren et al. 2014), and *ii*) physical forces between individuals that cause local mechanical instabilities at the interfaces between cell-types (Goldschmidt, Regoes et al. 2017, Borer, Ciccarese et al. 2020, Goldschmidt, Caduff et al. 2021).

Because metabolic interactions can direct the spatial positionings of different cell-types during the SSO process, we hypothesized here that they can also control the horizontal spread of functional novelty. Consider the conjugative transfer and proliferation of plasmid-encoded antibiotic resistance, which is of critical importance to human health. Plasmid conjugation typically requires direct contact between a plasmid donor and a potential recipient cell (Flemming and Wingender 2010, Stalder and Top 2016). We therefore expect that metabolic interactions that promote higher spatial intermixing of cell-types also promote increased plasmid conjugation, as there are more cell-cell contacts between cell-types (Fig. 1A). The ability of a plasmid to proliferate after successful conjugation requires that transconjugants (*i.e*., individuals of the potential recipient that received a plasmid) have access to growth-enabling resources (e.g., nutrients, unoccupied space, etc.). Because metabolic interactions can direct the spatial positionings of cell-types during range expansion (Goldschmidt, Regoes et al. 2017, Borer, Ciccarese et al. 2020, Goldschmidt, Caduff et al. 2021), we expect that metabolic interactions also determine the ability of transconjugants to proliferate (Fig. 1B). For example, if a metabolic interaction tends to position potential recipients behind the expansion frontier where resources are scarce, then transconjugants are less likely to proliferate (Fig. 1B). Conversely, if a metabolic interaction tends to position potential recipients at the expansion frontier where resources are plentiful, then transconjugants are more likely to proliferate (Fig. 1B).

**Figure 1.**
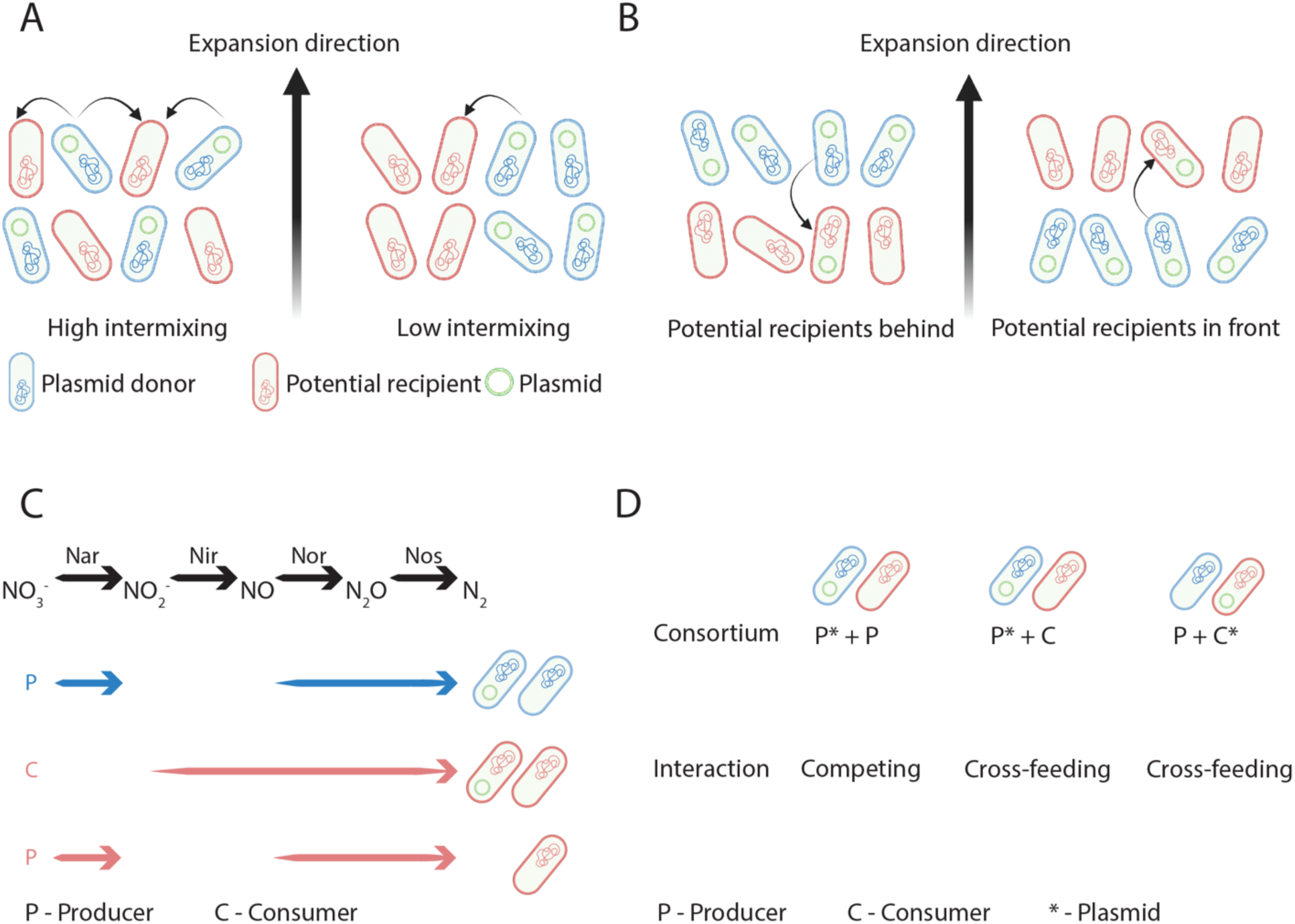
Hypothesis, expectations, and experimental system used in this study. We hypothesized that metabolic interactions control the spread of plasmid-encoded functional novelty. **A**, We expect that the level of spatial intermixing determines the extent of plasmid conjugation during range expansion. In this example, plasmid donors (blue cells) and potential recipients (red cells) undergo range expansion together with different levels of spatial intermixing. Curved arrows indicate potential plasmid conjugation events. High spatial intermixing results in a larger number of potential plasmid conjugation events. **B**, We further expect that the spatial positionings of potential recipients determine the ability of transconjugants to proliferate during range expansion. When transconjugants are positioned behind the expansion frontier where resources are scarce, they are less likely to proliferate. In contrast, when transconjugants are positioned at the expansion frontier where resources are plentiful, they are more likely to proliferate. **C**, We tested our hypothesis and expectations using five isogenic mutant strains of *P. stutzeri* A1601 that differ in their ability to reduce nitrogen oxides. Three strains have a loss-of-function deletion in the *nirS* gene and can reduce nitrate (NO_3_^-^) to nitrite (NO_2_^-^) but not nitrite (referred to as producers or P) while the other two have a loss-of-function deletion in the *narG* gene and can reduce nitrite to nitrogen gas (N_2_) but not nitrate (referred to as consumers or C). Each strain also carries a chromosomally-located gene encoding for cyan or red fluorescent protein (indicated by the colored arrows). Finally, each strain can carry plasmid pMA119, which encodes for kanamycin resistance and green fluorescent protein (indicated by the green circles). Definitions: Nar, nitrate reductase encoded by the *nar* genes; Nir, nitrite reductase encoded by the *nir* genes; Nor, nitric oxide reductase encoded by the *nor* genes; Nos, nitrous oxide reductase encoded by the *nos* genes. **D**, Our experimental design consists of three different consortia that undergo range expansion. P*+P; we imposed a competitive interaction between two producers, where each expresses a different chromosomally-encoded fluorescent protein and one carries pMA119. P*+C; we imposed a nitrite cross-feeding interaction between a producer and consumer, where each expresses a different chromosomally-encoded fluorescent protein and the producer carries pMA119. P+C*; we again imposed a nitrite cross-feeding interaction as described for P*+C, except in this case the consumer carries pMA119.

To test our hypothesis and expectations, we assembled pairs of strains of the bacterium *Pseudomonas stutzeri* into consortia where one strain was the plasmid donor and the other a potential recipient. We next imposed a competitive or resource cross-feeding interaction between the strains and conducted range expansion experiments, where we allowed the strains to expand and self-organize across space. We then combined image analyses of the experiments with spatially-explicit individual-based computational modelling to elucidate the causal pathway between metabolic interactions, spatial intermixing, spatial positionings, and the spread of a plasmid-encoded trait of critical importance to human health.

## Results

### Experimental system for quantifying plasmid conjugation and transconjugant proliferation during microbial range expansion

We constructed an experimental system to test our main hypothesis and expectations using consortia assembled from pairs of isogenic mutant strains of the facultative denitrifying bacterium *P. stutzeri* A1601 (Fig. 1C and Appendix – Table 1). One strain can reduce nitrate (NO_3_^-^) to nitrite (NO_2_^-^) but not nitrite to nitric oxide (NO) (referred to as the producer or P) while the other can reduce nitrite to nitric oxide but not nitrate to nitrite (referred to as the consumer or C) (Lilja and Johnson 2016) (Fig. 1C). Both strains carry a chromosomally-located gene that encodes for cyan (encoded by *ecfp*) or red (encoded by *echerry*) fluorescent protein (Goldschmidt, Regoes et al. 2017, Lilja and Johnson 2017) (Fig. 1C). In an anoxic environment with nitrate as the growth-limiting resource, two producers grown together engage in a competitive interaction for nitrogen oxides and other resources while the producer and consumer grown together engage in a nitrite cross-feeding interaction (Lilja and Johnson 2016, Goldschmidt, Regoes et al. 2017). For this study, we introduced the conjugative plasmid pMA119 into the producer or consumer (referred to as P* or C*, respectively) (Fig. 1C), where pMA119 carries genes encoding for kanamycin resistance and green fluorescent protein (Geisenberger, Ammendola et al. 1999). We could then use the expression of green fluorescent protein in conjunction with the expression of cyan or red fluorescent protein to identify the spatial locations and quantify the proliferation of pMA119 during range expansion. For our main experiments, we assembled different pairs of strains together into consortia to impose different metabolic interactions (Fig. 1D) and measured the consequences on *i*) spatial intermixing, *ii*) pMA119 conjugation, and *iii*) transconjugant proliferation during range expansion.

### Metabolic interactions determine the extent of plasmid pMA119 conjugation between strains during microbial range expansion

We first tested our expectation that metabolic interactions that promote higher levels of spatial intermixing between strains also promote greater extents of pMA119 conjugation (Fig. 1A). To accomplish this, we introduced pMA119 into one strain from each consortium (Fig. 1C,D) and demonstrated that pMA119 itself has no quantitative effect on the level of spatial intermixing that emerges during range expansion (Appendix – Figure 1). We then performed range expansion experiments in the absence of antibiotic selection for pMA119. After seven days of range expansion, we added kanamycin to the expansion areas and incubated the consortia for a further seven days. This imposed selection for transconjugants and allowed them to sufficiently proliferate such that we could experimentally identify and quantify individual pMA119 conjugation events (Fig. 2). As expected, we observed higher densities of plasmid pMA119 conjugation events (number of events per unit expansion area) when we imposed the nitrite (NO2^-^) cross-feeding interaction (P*+C and P+C*) than the competitive interaction (P*+P) (two-sample two-sided Welch test; *P_adj_* = 0.039, n = 5), regardless of whether the producer or consumer was the pMA119 donor (Fig. 2B). We verified that the transconjugants express the expected fluorescent proteins by plating on lysogeny broth (LB) agar plates amended with kanamycin (Appendix – Figure 3). We also performed color-swap experiments to confirm that our results are independent of the chromosomally-encoded fluorescent protein expressed by each strain (Appendix – Figure 4). We further compared the frequencies of transconjugants (area of transconjugants divided by the total expansion area) (Fig. 2C) and the mean areas of individual transconjugant regions (Fig. 2D). We found that both quantities are larger for the nitrite cross-feeding interaction than for the competitive interaction (two-sample two-sided Welch tests; *P_adj_* < 0.04, n = 5) (Fig. 2C,D). Thus, our data provide experimental evidence that metabolic interactions can indeed determine the number of plasmid conjugation events between different strains during range expansion.

**Figure 2.**
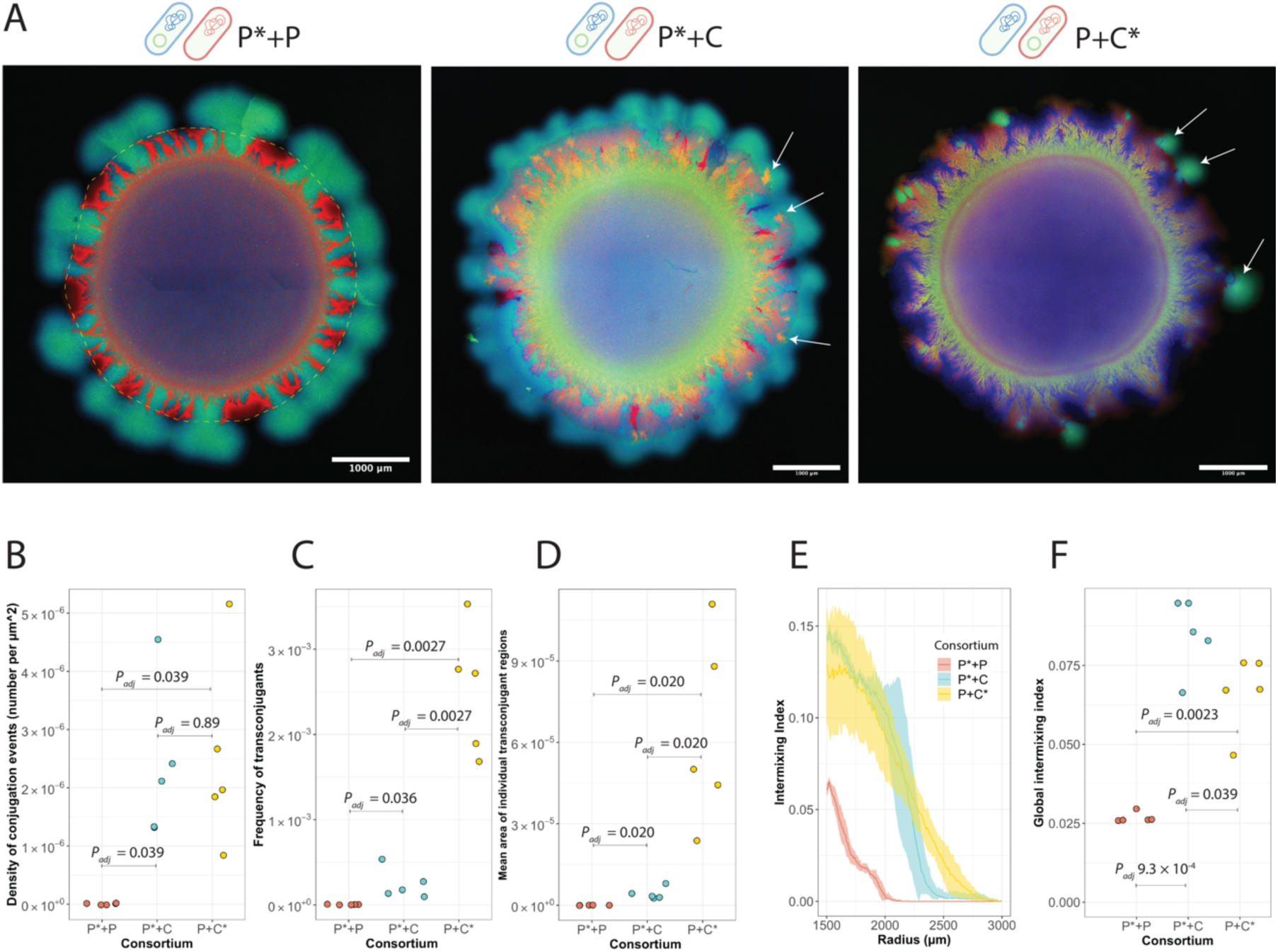
Metabolic interactions determine the extent of plasmid pMA119 conjugation during range expansion. **A**, The two schematic cells above each image indicate the two strains used for each experiment. Images are representative range expansions for consortia consisting of two producers that engaged in a competitive interaction where one producer carried pMA119 while the other did not (P*+P), a producer and consumer that engaged in a nitrite (NO_2_^-^) cross-feeding interaction where the producer carried pMA119 while the consumer did not (P*+C), and a producer and consumer that engaged in a nitrite cross-feeding interaction where the consumer carried pMA119 while the producer did not (P+C*). **B**, Density of pMA119 conjugation events (number of conjugation events per µm^2^ of expansion area) for each consortium. **C**, Frequency of transconjugants across the total expansion area (area of transconjugants divided by the total expansion area) for each consortium. **D**, Mean area of individual transconjugant regions for each consortium. **E**, Intermixing index as a function of radius from the edge of the inoculation area (1500 µm) to the edge of the final expansion frontier (3000 µm) at radial increments of 10 µm. Data are presented for independent experimental replicates (n = 5) and the shaded regions are the standard deviations at each radial increment. **F**, Global intermixing index measured as the sum of intermixing indices across the expansion area at radial increments of 10 µm. For **B-D**,**F**, each data point is the measurement for an independent experimental replicate (n = 5) and *P_adj_* is the Benjamini-Hochberg-adjusted *P* for a two-sided two-sample Welch test. P, producer; C, consumer; *, pMA119 carrier.

We next quantified and compared the levels of spatial intermixing across the expansion areas when imposing the competitive or nitrite (NO_2_^-^) cross-feeding interaction for the range expansion experiments where we added kanamycin after seven days. We observed higher levels of spatial intermixing for the nitrite cross-feeding interaction than for the competitive interaction regardless of whether the producer or consumer was the plasmid pMA119 donor (Fig. 2E,F). This is also clear from high-resolution images, where the competitive interaction resulted in nearly complete spatial segregation of the strains while the nitrite cross-feeding interaction resulted in high spatial intermixing at the level of individual cells (Appendix – Figure 5). We further validated that the conjugation efficiency of pMA119 is identical for the producer and consumer and independent of the metabolic interaction that we imposed (Appendix – Figures 6-8). Thus, adding kanamycin does not alter our general conclusion that nitrite cross-feeding promotes higher levels of spatial intermixing during range expansion.

### Spatial intermixing determines the extent of plasmid pMA119 conjugation between cell-types during microbial range expansion

We next tested whether spatial intermixing itself determines the extent of plasmid conjugation during range expansion. To accomplish this, we modified and adapted a previously described spatially-explicit individual-based computational model (Rudge, Steiner et al. 2012). See the method section for a full documentation of our implementation of the model. Briefly, we modelled individual bacterial cells as three-dimensional capsules that grow, divide, and interact according to user-specified rules. We first validated the utility of the model to manipulate spatial intermixing. To accomplish this, we imposed three levels of initial spatial intermixing between two cell-types, where one cell-type is a plasmid donor (cyan cells) and the other a potential recipient (red cells). The three levels of spatial intermixing are “low spatial intermixing” where we initially spatially segregated competing cell-types, “intermediate spatial intermixing” where we initially positioned competing cell-types according to a checkerboard arrangement, and “high spatial intermixing” where we initially positioned cross-feeding cell-types according to the same checkerboard arrangement. For each simulation, we positioned the cells across a surface with a uniform distance between individuals and random rotational orientation of individuals along a two-dimensional plane (Fig. 3A). Competing cells have independent growth rates whereas cross-feeding cells have growth rates that are dependent on a diffusible signal that emerges from their cross-feeding partner cells. If a plasmid donor cell comes into physical contact with a recipient cell, we applied a constant probability of plasmid conjugation, whereupon the recipient cell would become a transconjugant (yellow cells).

**Figure 3.**
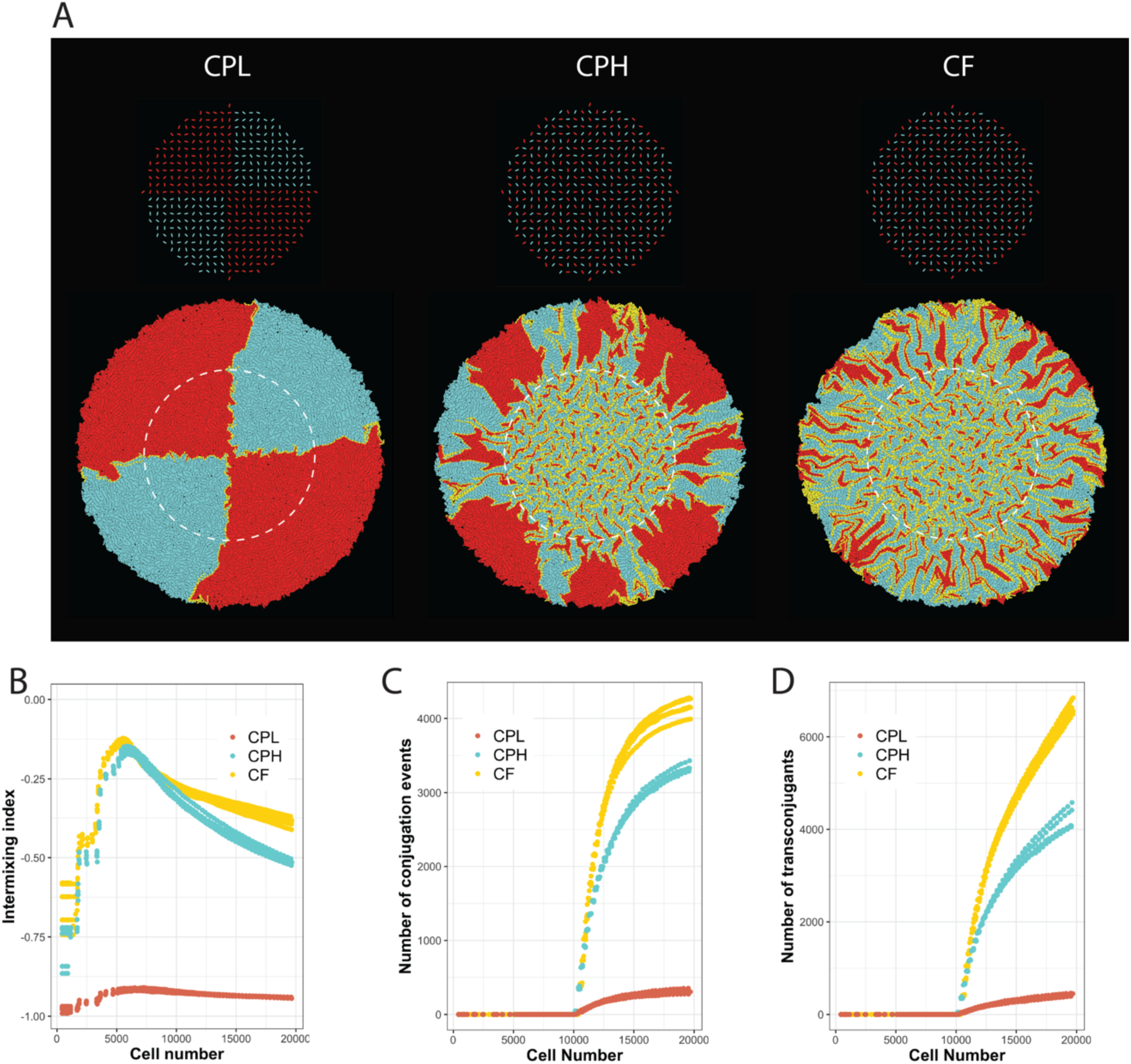
Spatial intermixing determines the extent of plasmid conjugation during range expansion. **A**, Images are representative individual-based computational simulations of range expansions for low, intermediate and high spatial intermixing at the end of the simulations. We simulated the growth of two cell-types that engage in a competitive or cross-feeding interaction, where one is a plasmid donor and the other a potential recipient. CPL; competitive interaction with low spatial intermixing. CPH; competitive interaction with high spatial intermixing. CF; cross-feeding interaction with high spatial intermixing. The initial positions of individual cells are displayed above and the final expansions below. White dashed circles indicate the inoculation area. Cyan cells are plasmid donors, red cells are potential recipients, and yellow cells are transconjugants. **B**, Intermixing index as a function of population size (cell number). **C**, Accumulated number of plasmid conjugation events during range expansion as a function of population size (cell number). **D**, Accumulated number of transconjugants during range expansion as a function of population size (cell number). For **B-D**, cell number is comparable to time as cell number increased monotonically with time. Plasmid conjugation was only possible after the total cell number reached 10,000. Data are presented for five independent simulations for each interaction and for each level of initial spatial intermixing.

We found that the initial level of spatial intermixing determined the final level of spatial intermixing of our simulations; the final level of spatial intermixing for the competing cell-types was significantly higher when the initial level of spatial intermixing was higher (two-sample two-sided Welch tests; *P_adj_* = 3.9 x 10^-8^, n = 5) (Fig. 3B). We further found that the cross-feeding interaction resulted in higher spatial intermixing than the competitive interaction (two-sample two-sided Welch tests; *P_adj_* < 2.8 x 10^-6^, n = 5) (Fig. 3B), which is consistent with our experimental results (Fig. 2). In our simulations, cells were initially not in physical contact with other cells (Fig. 3A), and thus grew into microcolonies that filled the available space. This is the cause for the initial rise in the intermixing index (Fig. 3B). As cells subsequently grew outwards in the radial direction, the intermixing index declined as we observed in our experiments (Fig. 2). The emergent patterns for the simulations with the initial checkerboard (Fig. 3A) and those from our experiments (Fig. 2) are qualitatively similar, indicating that our computational model successfully incorporates key features of the SSO process.

We then analyzed the effect of spatial intermixing on plasmid conjugation by tracking the number of conjugation events (Fig. 3C) and the number of transconjugants (Fig. 3D). For our simulations, we did not allow conjugation to occur until we observed clear deviations in the temporal dynamics of the intermixing index among the different consortia, which occurred after the population sizes exceeded approximately 10,000 cells (note the point at which the yellow and blue datapoints in Fig. 3B begin to deviate). This allowed us to isolate the effects of intermixing on conjugation. For the competing interaction, we found that both the numbers of conjugation events and the numbers of transconjugants were larger when we imposed high spatial intermixing (two-sample two-sided Welch test; *P_adj_* < 3.9 x 10^-8^, n = 5) (Fig. 3C,D). We also found that both the numbers of conjugation events and the numbers of transconjugant cells were larger when we imposed the cross-feeding interaction rather than the competitive interaction (two-sample two-sided Welch test; *P_adj_* < 9.4 x 10^-6^, n = 5) (Fig. 3C,D). Thus, spatial intermixing does indeed have direct effects on plasmid conjugation between different cell-types during range expansion, as it determines the number of cell-cell contacts and thus the number of opportunities for plasmid conjugation.

### Spatial positionings of potential plasmid recipients determine the proliferation of transconjugants during microbial range expansion

We next tested our second expectation that the spatial positionings of potential plasmid recipients determine the ability of transconjugants to proliferate during range expansion (Fig. 1B). For our nitrite (NO_2_^-^) cross-feeding system, the expansion frontier is predominantly occupied by the producer where resources are readily available (Goldschmidt, Regoes et al. 2017, Goldschmidt, Caduff et al. 2021). We therefore expected transconjugants of the producer to proliferate more extensively than transconjugants of the consumer. Indeed, we found that the mean areas of individual transconjugant regions were significantly larger when the producer was the potential recipient than when the consumer was (two-sample two-sided Welch test; *P_adj_* = 0.020, n = 5) (Fig. 2D).

To gain further evidence that spatial positionings determine the ability of transconjugants to proliferate, we performed additional experiments where we set the initial inoculum ratio of the producer-to-consumer for the nitrite (NO_2_^-^) cross-feeding interaction to 1 or 1000. The initial ratio determines the initial extent to which the producer or consumer occupy the expansion frontier(Goldschmidt, Caduff et al. 2021), and we therefore reasoned it would also determine the ability of transconjugants to proliferate. When the ratio is 1, each position at the initial expansion frontier is randomly occupied by a producer or consumer (referred to as “Random frontier”) (Fig. 4A, D). When the ratio is 1000, the producer is more likely to occupy each position at the initial expansion frontier (referred to as “Producer frontier”) (Fig. 4A, D). To complement our experiments, we again performed individual-based computational simulations with two cross-feeding cell-types as described above, but in this case, we varied the initial spatial positionings of the producer and consumer while fixing the initial ratio of producer-to-consumer to 1:1. We deposited 219 plasmid donor and 218 potential recipient cells across a surface with two different initial spatial distributions of the producer and consumer at the expansion frontier. These two initial spatial distributions are “Random frontier” where we randomly designated each cell at the initial expansion frontier as a producer or consumer, and “producer frontier” where we designated each cell at the initial expansion frontier as a producer (Appendix – Figure 9). We found that, for both the experiments and simulations regardless of whether the producer or consumer was the pMA119 recipient, the numbers of pMA119 conjugation events that proliferated to an experimentally observable extent were significantly lower when more producers were initially positioned at the expansion frontier (two-sample two-sided Welch test; *P* < 0.05, n = 5) (Fig. 4C, F). This is because, as the producer increases in dominance at the initial expansion frontier, transconjugants occur further behind the frontier where resources are scarce and transconjugants are less likely to proliferate. We colored individual transconjugant lineages differently in order to track their development during range expansion (Fig. 4B, E). One color represents one unique lineage. We found that lineage size decreased as more producers were initially positioned at the expansion frontier (Appendix – Figure 10A). We also quantified the frequencies of transconjugants as we did for Fig. 2, and both quantities showed significantly lower values when more producers were initially positioned at the expansion frontier (Appendix – Figures 10B, C). Thus, consistencies between experiments and modelling simulations provide further evidence that spatial positioning is an important determinant of transconjugant proliferation.

**Figure 4.**
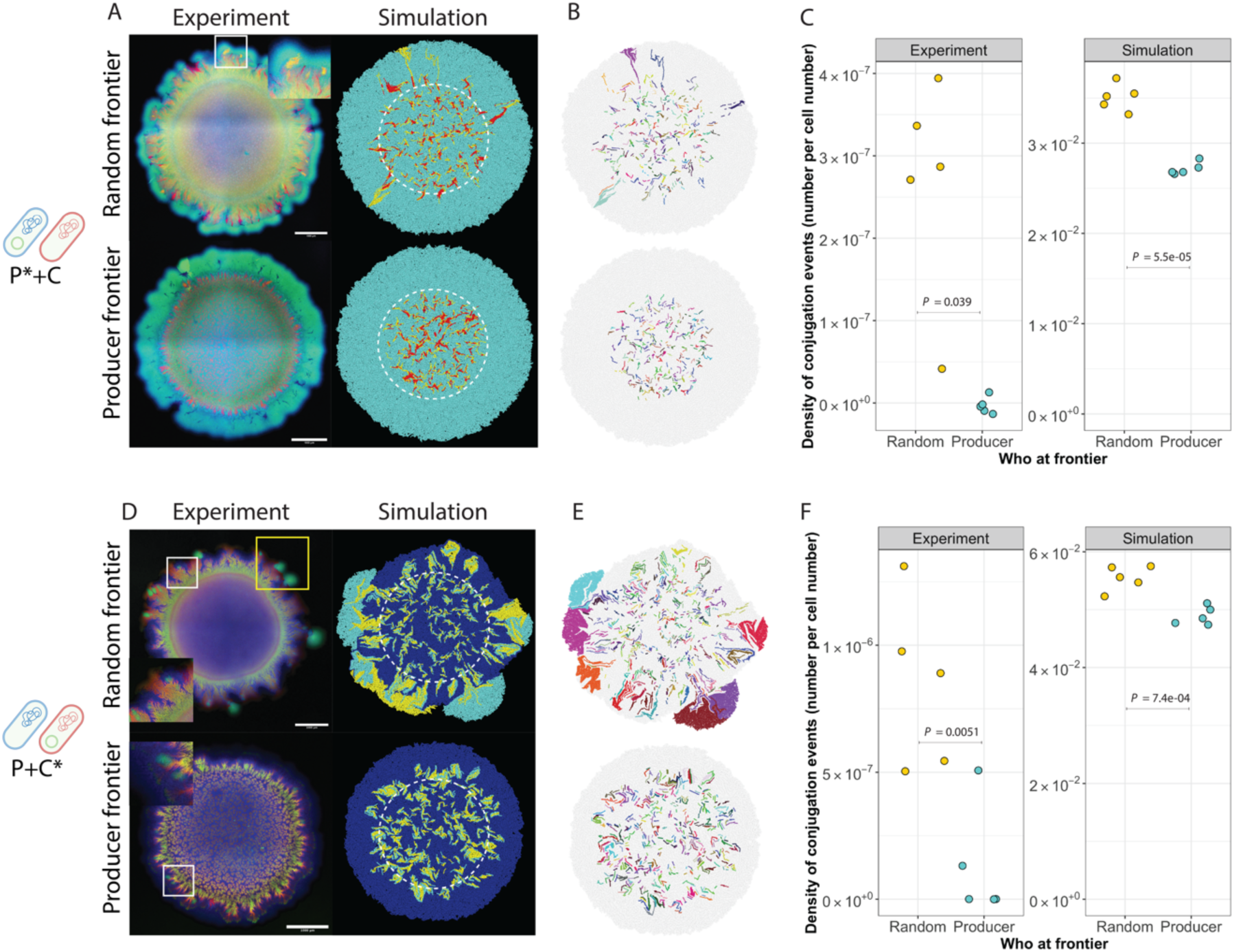
Effect of spatial positioning on the extent of plasmid pMA119 conjugation and transconjugant proliferation during range expansion. The two schematic cells on the left side of the image indicate the two strains used for each experiment. Images are representative range expansions for consortia consisting of a producer and consumer that engaged in a nitrite (NO_2_^-^) cross-feeding interaction where the producer carried pMA119 while the consumer did not (P*+C), and a producer and consumer that engaged in a nitrite cross-feeding interaction where the consumer carried pMA119 while the producer did not (P+C*). We experimentally set the initial ratio of producer-to-consumer to 1 (Random frontier) or 1000 (Producer frontier).For the simulations, we kept the ratio of producer-to-consumer to 1 but manipulated initial positioning. **A**, Consortia consisting of P*+C. Transconjugants are colored yellow and the white frame indicates observable transconjugant regions. Left panels are experiment results, and right panels are simulation results. **D**, Consortia consisting of P+C*. Transconjugants are colored cyan. White frames indicate observable transconjugant regions and yellow frames incidate transconjugant regions that appear at the expansion frontier. Left panels are experiment results and right panels are simulation results. **B, E**, Corresponding transconjugant lineages of the simulations in **A** and **D**. Each individual lineage is labelled with a different color. **C, F**, Density of conjugation events (number of conjugation events divided by the total number of cells) for each initial expansion frontier.

## Discussion

Using a combination of experiments and spatially-explicit individual-based computational simulations, we demonstrated that spatial intermixing determines the number of conjugation-mediated plasmid transfer events during range expansion (Figs. 2 and 3). While several studies have previously reported that metabolic interactions can determine the level of spatial intermixing during range expansion (Momeni, Brileya et al. 2013, Müller, Neugeboren et al. 2014, Goldschmidt, Regoes et al. 2017), the connection between spatial intermixing and plasmid conjugation has remained unclear. For example, while recent studies demonstrated that hydrodynamic conditions that increase the number of cell-cell contacts can also promote plasmid conjugation (Tecon, Ebrahimi et al. 2018, Ruan, Ramoneda et al. 2021), these studies did not explicitly consider the local spatial arrangements of different cell-types. Thus, an important outcome of our study is the establishment of a causal pathway between metabolic interactions, local spatial intermixing, and conjugation-mediated plasmid transfer.

We further provided novel insights into the mechanisms that determine the ability of transconjugants to proliferate during range expansion. Using experiments and computational modelling, we showed that metabolic interactions direct the spatial positionings of plasmid donors and potential recipients, which in turn determine the ability of transconjugants to proliferate (Fig. 4). In our case, transconjugants proliferated more effectively when more consumers were positioned at the initial expansion frontier, regardless of whether the consumer or producer was the plasmid donor (Fig. 4). Why is this? Consider a scenario where the consumer is the potential recipient and the producer is the plasmid donor. If the consumer occupies the initial expansion frontier, then consumer transconjugants can emerge at the expansion frontier where resources and unoccupied space are plentiful, which will promote the proliferation of transconjugants (Appendix – Figure 11A). Conversely, if the producer occupies the initial expansion frontier, then consumer transconjugants will emerge behind the expansion frontier where resources and unoccupied space are scarce, which will repress transconjugant proliferation (Appendix – Figure 11B). Now consider a scenario where the producer is the potential recipient and the consumer is the plasmid donor. If the consumer occupies the initial expansion frontier, then producer transconjugants can emerge close to the expansion frontier where resources and unoccupied space are plentiful, which again will promote transconjugant proliferation (Appendix – Figure 11C). Conversely, if the producer occupies the initial expansion frontier, then the consumer will lie behind the frontier. Producer transconjugants will therefore also only emerge behind the expansion frontier where resources and unoccupied space are scarce, which again will repress transconjugant proliferation (Appendix – Figure 11D). Thus, initial spatial positioning in conjunction with the interaction-dependent dynamic process of SSO determine whether transconjugants can proliferate.

Plasmid conjugation is a process that generates functional novelty to microbial communities, but this process has not been investigated in detail during range expansion. In some ways, understanding the fate of transconjugants has parallels with understanding the fate of random genetic mutations. The fate of random genetic mutations is fairly well-understood (Hallatschek, Hersen et al. 2007, Hallatschek and Nelson 2010, Gralka, Stiewe et al. 2016, Bosshard, Dupanloup et al. 2017, Goldschmidt, Regoes et al. 2017, Bosshard, Peischl et al. 2019, Gralka and Hallatschek 2019, Yu, Gralka et al. 2021). For example, both transconjugants and random genetic mutations are susceptible to selection and strong drift at the expansion frontier. Nevertheless, there are also fundamental differences. One important difference is the origin of the genetic change. As opposed to random genetic mutations, plasmid conjugation typically requires direct cell-cell contact(Sørensen, Bailey et al. 2005, Thomas and Nielsen 2005, Babic, Lindner et al. 2008, Hayes, Aoki et al. 2010, Stalder and Top 2016) and therefore depends on the spatial organization and positionings of different cell-types (Fig. 2). These spatial positionings, however, are context dependent (e.g., they depend on the types of metabolic interactions that occur between different cell-types) and dynamic (Fig. 2). Thus, predicting the origin and subsequent fate of transconjugants requires additional information on the underlying mechanisms driving the SSO process.

Our main conclusions are potentially generalizable to any surface-associated microbial community and to any plasmid that is transferred via conjugation, including communities and plasmids important for human health and disease, environmental remediation, and biotechnology. Our first conclusion, which is that the extent of spatial intermixing of different cell-types determines the extent of plasmid conjugation, is based on a fundamental rule; plasmid conjugation requires cell-cell contact between a plasmid donor and a potential recipient. Increased spatial intermixing results in increased numbers of cell-cell contacts, and thus increased possibilities for plasmid conjugation events. Because of this fundamental rule, we believe our conclusion will be generally valid for any plasmid that requires cell-cell contact for conjugation, regardless of what traits are conferred by the plasmid. Our second conclusion, which is that the proliferation of transconjugants depends on spatial positioning, is based on a fundamental constraint; transconjugants cannot proliferate unless they are positioned such that they have access to growth-supporting resources. We therefore believe our main conclusions will be of broad relevance to nearly any microbial system where plasmid-encoded traits are important.

## Materials and Methods

### Bacterial strains and plasmid

We reported a detailed description of the construction of the *P. stutzeri* strains used in this study elsewhere (Lilja and Johnson 2016, Goldschmidt, Regoes et al. 2017) and provide the genotypes and phenotypes of the strains in Appendix – Table 1. Briefly, the producer has a loss-of-function deletion in the *nirS* gene and cannot reduce nitrite (NO_2_^-^) to nitric oxide (NO) while the consumer has a loss-of-function deletion in the *narG* gene and cannot reduce nitrate (NO_3_^-^) to nitrite. Note that both strains have an intact periplasmic nitrate reductase encoded by the *nar* genes, but this reductase does not support growth of *P. stutzeri* under anoxic conditions (Lilja and Johnson 2016, Goldschmidt, Caduff et al. 2021). The producer carries either an isopropyl ß-D-1-thiogalactopyranoside (IPTG)-inducible cyan (*ecfp*) or red (*echerry*) fluorescent protein-encoding gene while the consumer only carries the *echerry* gene. We introduced both fluorescent protein-encoding genes into the same neutral site located in the chromosome using derivatives of the mini-Tn7T transposon as we reported elsewhere (Lilja and Johnson 2016, Goldschmidt, Regoes et al. 2017). For this study, we further introduced the GFPmut3b-tagged plasmid pMA119 into each of the strains, which is a derivative of the broad host-range plasmid RP4(Geisenberger, Ammendola et al. 1999). Briefly, the pMA119 plasmid encodes for an IPTG-inducible variant of green fluorescent protein and confers kanamycin resistance.

### Construction of plasmid donor strains

To import plasmid pMA119 into the producer or consumer and create the experimental plasmid donor strains, we first grew *Pseudomonas putida* SM1443 carrying pMA119 overnight in liquid LB medium amended with 50 µg mL^-1^ kanamycin to prevent the proliferation of plasmid segregants. We additionally grew the producer and consumer separately overnight in liquid LB medium. We then independently adjusted the OD_600_ of each overnight culture to 2.0 with 0.89% (w/v) sodium chloride solution, mixed *P. putida* SM1443 with either the producer or consumer at a volumetric ratio of 1:1, deposited a 1 µL aliquot from each mixture onto the surface of a separate LB agar plate, and incubated the droplet for 24h at 30°C. After incubation, we removed the resulting colonies using sterile inoculation loops and suspended the biomass in 0.89% (w/v) sodium chloride solution. We then serially diluted the cell suspensions and plated each dilution onto LB agar plates containing kanamycin (50 µg mL^-1^) to select for *P. stutzeri* transconjugants, gentamycin (10 µg mL^-1^) to select against *P. putida* SM1443, and IPTG (0.1mM L^-1^) to verify transconjugants. Successful delivery of pMA119 results in the producer or consumer that is resistant to both kanamycin and gentamycin, expresses green fluorescent protein (from pMA119), and additionally expresses cyan or red fluorescent protein (from the chromosome).

### Range expansion experiments

We used a modified version of a range expansion protocol described in detail elsewhere(Goldschmidt, Regoes et al. 2017). Briefly, we grew the plasmid donor strain overnight with LB medium amended with kanamycin (50 µg mL^-1^) to maintain plasmid pMA119 and the recipient strain overnight in LB medium without antibiotics. We then independently adjusted the OD_600_ of each overnight culture to 2.0 with 0.89% (w/v) sodium chloride solution, mixed one plasmid donor and one potential recipient culture together at a volumetric ratio of 1:1 as indicated in the results section, and transported the mixtures into a glove box (Coy Laboratory Products, Grass Lake, MI) containing an anoxic nitrogen (N_2_):hydrogen (H_2_) (97%:3%) atmosphere. We next deposited 1 µL aliquots of the culture mixture onto the surfaces of separate LB agar plates amended with 0.1 mM IPTG and 1 mM nitrate (NO_3_^-^) as the growth-limiting substrate and incubated the LB agar plates at 21 °C for two weeks inside the glovebox. For experiments where we added kanamycin, we removed the LB agar plates from the incubator (located inside the glove box) after one week of expansion, deposited 10 µL of kanamycin (final concentration in the LB agar plate of 50 µg mL^-1^) to the LB agar plates at four points located approximately 7 mm away from the expansion center using a self-made mold (Appendix – Figure 12), and then returned the LB agar plates to the incubator for a second week of expansion. After two weeks of incubation, we removed the LB agar plates from the glove box and incubated them at 4°C overnight in ambient air to promote maturation of the fluorescent proteins. The patterns of spatial self-organization and expansion areas did not significantly change during incubation at 4°C in ambient air. We performed five biological replicates for all of our experiments.

### Color-swap experiment

We performed color-swap experiments to test for any effects of the chromosomally-encoded fluorescent protein-encoding gene on pMA119 conjugation using the same range-expansion experimental protocols as described in the Methods section. The only difference is that we used a producer strain carrying a chromosomally-located echerry gene (encoding for red fluorescent protein) and a consumer strain carrying a chromosomally-located *ecfp* gene (encoding for blue fluorescent protein). Note that for our main experiments we used a producer strain carrying a chromosomally-located *ecfp* gene and a consumer strain carrying a chromosomally-located *echerry* gene (Fig. 1C,D). We reported the data in Appendix – Figure 2.

### Full spectrum flow cytometry and data analysis

We obtained all the conjugation rate measurements reported in Appendix – Figures 3 and 4 and Fig. 4 using a Cytek Aurora spectral flow cytometer (Cytek Biosciences, Amsterdam, NL) operated by the Flow Cytometry Core Facility at ETH Zürich (https://facs.ethz.ch). This flow cytometer contains a full-spectrum analyzer that can capture a wide array of fluorochrome combinations. We performed data post-processing and presentation using FlowJo 10.8.0 software (https://www.flowjo.com).

### Preparation of control cultures for full-spectrum flow cytometry

Before we performed full-spectrum flow cytometry for our experiments, we first prepared controls in order to establish settings to detect and discriminate the different fluorescent proteins that we used in our experiments. Our controls consisted of the following four strains; *Pseudomonas stutzeri* A1603 that expresses cyan fluorescent protein (Appendix – Table 1), *P. stutzeri* A1603 that expresses red fluorescent protein (Appendix – Table 1), *Pseudomonas putida* SM1443 carrying plasmid pMA119 (RP4::gfpmut3b) that expresses green fluorescent protein, and *P. stutzeri* A1501 that does not express any fluorescent proteins. The first three strains provided single fluorochrome information while the fourth strain provided background information on cell size and served as a negative control. We grew all of the strains individually overnight in lysogeny broth (LB) medium with shaking at 37°C. For *P. putida* SM1443, we amended the LB medium with 50 ug ml^-1^ kanamycin to prevent plasmid segregants from proliferating. After overnight growth, we washed the cells with phosphate-buffered saline (PBS) prior to flow cytometric analysis.

### Sample preparation for quantifying plasmid pMA119 conjugation using filter mating

We quantified pMA119 conjugation using a conventional filter mating approach (Appendix – Figure 3) with three pairs of donor and recipient strains (P*+P, P*+C, and P+C*) (see Fig. 1d for a description of the composition of each pair). We first grew each strain individually overnight with oxic LB medium in a shaking incubator at 37°C. We then adjusted the OD_600_ of each overnight culture to 2 with 0.89% (w/v) sodium chloride solution. We next mixed the individual cultures into pairs at volumetric ratios of 1:1, inoculated 50 µL aliquots of each mixture onto a separate filter (pore size of 0.22 µm) lying directly on the surface of an LB agar plate, and incubated the LB agar plates for 24 hours at 37°C. After incubation, we washed cells off the filters using phosphate-buffered saline and prepared the cells for full-spectrum flow cytometry. We performed this experiment with three experimental replicates.

### Sample preparation for quantifying plasmid pMA119 conjugation in anoxic batch culture

We also quantified pMA119 conjugation in anoxic batch cultures (Appendix – Figure 4) using the same three pairs of donor and recipient strains (P*+P, P*+C, and P+C*) (see Fig. 1d for a description of the composition of each pair). Our objective of this experiment was to test whether the metabolic interaction imposed between the strains (competition or nitrite [NO_2_^-^] cross-feeding) modifies the conjugation efficiency of pMA119. We first prepared batch cultures containing 10 mL anoxic LB medium with 3 mM nitrate (NO_3_^-^) in a glove box containing an anoxic nitrogen (N_2_):hydrogen (H_2_) (97%:3%) atmosphere as described in the Methods section. We adjusted the pH of the batch cultures to pH 7.5 to prevent nitrite toxicity (Sijbesma, Almeida et al. 1996, Zhou, Oehmen et al. 2011). We then grew each strain individually overnight with oxic LB medium in a shaking incubator at 37°C, adjusted the OD_600_ of each overnight culture to 2 with 0.89% (w/v) sodium chloride solution, mixed the overnight cultures into pairs at volumetric ratios of 1:1, and inoculated 1 µl aliquots of the mixtures into fresh anoxic LB medium. We then incubated the pairs overnight in a shaking incubator at 37°C and collected samples using syringes for full-spectrum flow cytometry. We performed this experiment with three experimental replicates.

### Image acquisition and analysis

We acquired confocal laser scanning microscope (CLSM) images of our range expansions using a Leica TCS SP5 II confocal microscope (Leica Microsystems, Wetzlar, Germany). We used the following objectives; 5x/0.12na (dry), 10x/0.3na (dry) and 63 x/1.4na (oil) (Etzlar, Germany). We used a frame size of 1024 x 1024 and a pixel size of 3.027 µm. We set the laser emission to 458 nm for the excitation of cyan fluorescent protein, to 488 nm for the excitation of green fluorescent protein, and to 514 nm for the excitation of red fluorescent protein.

We performed image analysis in ImageJ (https://imagej.nih.gov/ij/) with Fiji plugins (v. 2.1.0/1.53c) (https://fiji.sc). We first auto-thresholded channel four to obtain outlines of the expansion areas using the ‘Otsu dark’ function. We then quantified the areas and numbers of transconjugant regions using the Ilastik took kit (v. 1.3.3) (https://www.ilastik.org). Briefly, we first trained each image individually and exported the images back into Fiji as HDF5 files. We next applied the ‘Huang dark’ function to auto-threshold the HDF5 files and extract transconjugant regions, filtered out objects smaller than 100 square pixels that might result from noise (note that our criteria for identifying transconjugant regions was that regions must be sufficiently large to be experimentally observable at the community level), and applied the ‘analyze particle’ function to obtain counts and areas of transconjugant regions.

### Quantification of intermixing

We quantified spatial intermixing (referred to as the intermixing index) between strains from the CLSM images using Fiji (*v1.53c*) plugins (https://fiji.sc) as described in detail elsewhere(Goldschmidt, Regoes et al. 2017). Briefly, we first cropped all the images to squares and applied the ‘Otsu dark’ function to auto-threshold and the ‘Niblack’ function to auto local threshold channel five (red fluorescent protein). We then used the Sholl analysis plugin (Ferreira, Blackman et al. 2014) on channel five to calculate the number of intersections between background and information-containing parts of the image at 10 µm increments from the centroid of the range expansion to the outer edge of the expansion frontier. We next extracted the data between radii of 1500 and 3000 µm. This region excludes the inoculation area and captures the expansion region. We excluded radii less than 1500 µm for two reasons. First, they do not accurately capture the spatial features caused by the range expansion process. Second, fluorescent signals at smaller radii are difficult to precisely resolve, thus creating noise. To quantify the global intermixing index, we summed the individual intermixing indices at 10 µm radial increments between radii of 2000 and 3000 µm (Fig. 2) or between 1500 and 3000 µm (Fig. 3) and then normalized the sum by the number of radial increments that contained non-zero values. We selected these radii because they contain the points at which the intermixing index begins to decline (at 2000 µm in Fig. 2 and at 1500 µm in Fig. 3).

### Validation of transconjugants

We performed additional experiments to validate that the acquisition of green fluorescent protein accurately indicates plasmid pMA119 conjugation to potential recipients. To achieve this, we sampled individual transconjugant regions from the same range expansion experiments described above using sterile toothpicks and streaked them onto new LB agar plates amended with kanamycin (50 µg mL^-1^) (Appendix – Figure 3). We also validated that the fluorescence itself has no effect on pMA119 conjugation by swapping colors between plasmid donors and recipients (Appendix – Figure 4).

### Individual-based computational modelling

We built a spatially-explicit individual-based computational model to mimic our experimental system using the CellModeller 4.3 framework (Rudge, Steiner et al. 2012) (https://haselofflab.github.io/CellModeller/.), which is a python-based, open-source framework for modelling bacterial populations. In our implementation, we modelled individual rod-shaped bacterial cells as three-dimensional capsules (i.e., cylinders with hemispherical ends), which grow by extending their length. Capsules experience viscous drag and cannot grow into one another. As they grow, cells add a constant volume until they reach a critical size where they then divide into two daughter cells, ensuring cell size homeostasis. In CellModeller, cells are abstracted as computational objects referred to as a cellState (cs) that contains all the information regarding an individual cell, including its spatial position (pos[x, y, z]), rotational orientation (dir[x, y, z]), cell length (len), growth rate (growthRate), and cell type (cellType). The cell-type is an arbitrary label that allows us to simulate different cellular behaviors.

CellModeller also contains an integrator module that solves differential equations to model intracellular chemical dynamics, as well as a signaling module responsible for the diffusion of certain molecules in and out of cells and around the extracellular space. In simulations with a metabolic cross-feeding interaction, we modelled this by having the producer strain constantly producing a diffusing signal (named *A*). Although signal *A* is made inside producer cells, it also diffuses outside of producer cells and into the extracellular grid. For each consumer cell, we set the growth rate to be proportional to the intracellular concentration (*A_in_*) that had diffused in according to the following equation:

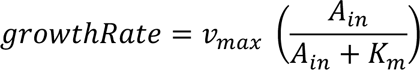

where v_max_ is the maximum consumer growth rate (set to 0.9) and K_m_ is the sensitivity of growth to signal A (set to 0.1). The producer *growthRate* was always set to 1.0. Note that in CellModeller other cells physically constrain the growth of any individual cell, so the *growthRate* parameter is a target amount of length added by each cell. When cells cannot exert sufficient force to push their neighbors and thus add length, their effective growth rate declines.

We extended our model to include plasmid conjugation. As part of the biophysics in CellModeller, physical contacts between cells are calculated at each step to minimize any overlap between cells. We altered the code such that each cell kept track of their contacts, and this allowed us to model plasmid transfer when cells were in contact. This function is activated by setting the argument ‘compNeighbours=True’ when initiating the biophysical model. When donor and recipient cells were in contact, we assumed a constant probability per unit time of plasmid transfer, set to Pc = 0.0005. Note that in order to better observe the plasmid conjugation process between populations, we only allowed plasmid transfer to occur between plasmid donor and potential recipient cells but not between transconjugants and potential recipient cells.

Given our interest in range expansion, we chose to focus on investigating the dynamics of conjugation primarily in the expansion area where the range expansion process occurs. To achieve this, we did not allow conjugation to occur until we observed clear deviations in the temporal dynamics of intermixing between different consortia and initial conditions (cell number > 10,000) (see the Appendix). As such, the simulated dynamics are mainly governed by the range expansion process, which reduces noise that would otherwise be generated from the inoculation area. When we did enable conjugation during the initial phases, we observed the emergence of transconjugant lineages that block further contacts between donor and recipient cells, thus obscuring the investigation of conjugation during the range expansion process itself.

#### Cell division

In CellModeller simulations, individual cells are modelled as cylinders of length l capped with hemispheres that result in a capsule shape, with both hemispheres and the cylinder having a radius r. At each simulation step, a cell increases in length based on its growth rate parameter, which is physically constrained by the other cells in its physical proximity. In this work, we initiated cells to have r = 0.04 and l = 2 and set cells to divide when their length reaches the critical division length l_&’(_ with the following equation:

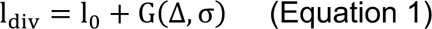

where l_)_ is the initial cell length at birth and G is a random gaussian distribution with mean Δ = 2 and standard deviation σ = 0.45. Therefore, when a cell divides, the two daughter cells are initiated with l_&’(_ /2 and a new target division length is assigned to each daughter cell calculated from Equation 1 above. The addition of constant mass has been found to accurately model bacterial division while maintaining cell size homeostasis as described elsewhere (Taheri-Araghi, Bradde et al. 2015).

#### Metabolic interactions

In simulations that include metabolic cross-feeding, we modeled a signal (referred to as signal A) that is produced inside cells and can diffuse in and out of neighboring cells and around the extracellular space. We modelled the extracellular space as a 3-dimensional grid with dimensions of 120 x 120 x 12, where each grid voxel has a size of 4 x 4 x 4. We calculated the concentration of A within a cell (A_’*_) with the following equation:

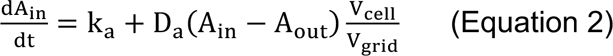

where k_-_is the constant production rate, D_-_is the diffusion constant across the cell membrane, A_./,_ is the concentration of A in the local grid voxel, V_1233_ is the volume of the cell, and V_45’&_ is the volume of the grid. For the consumer cells, the k_-_term was absent, so A_’*_ can only enter by diffusion. We determined the growth rates of consumer cells with the following equation:

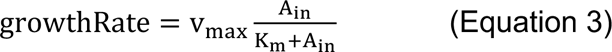

v_6-7_ is the maximal growth rate (set to 0.9). K_6_ determines the sensitivity of growth to signal A (set to 0.1). For comparison, the growth rate of the producer is always 1.

#### Initial cell layout

To initiate the simulations, we loaded 437 cells (219 donor cells and 218 recipient cells) across the grid with a uniform distance of 5 units between cells along the x and y axes, but only at grid points that were within a circle of radius 60 units from the origin. We loaded all the cells with the z coordinate = 0. Thus, we constrained their orientations and dynamics to the x,y plane. We manipulated the spatial positionings of cells by controlling the spatial distributions of cell-types across the grid. To obtain an expansion frontier completely occupied by the producer, we labeled the cells that were furthest from origin as producers and then labeled all the cells behind the fronter randomly. To obtain a frontier randomly occupied by the producer or consumer, we randomly labeled all cells.

#### Quantification of the intermixing index

To quantify the intermixing index (referred to as intermixing index II) from the simulations, we used the following equation:

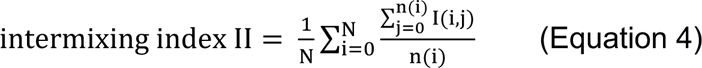

where N is the total number of cells and n(i) is the number of neighbors surrounding cell i. I(i, j) is either 1 or −1 depending on whether the neighbor has a cell-type that is different from (1) or the same as (−1) the focal cell. For an individual cell i, we iterated across all neighbors that are in physical contact j:

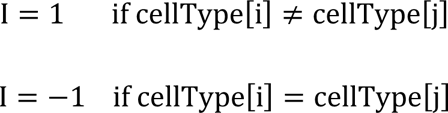

This method is similar to the spatial assortment parameter(Yanni, Márquez-Zacarías et al. 2019) or segregation index(Nadell, Foster et al. 2010) used in previous studies. The intermixing index II represents the degree of mixing among nearest neighbors across space. Areas with large domains of isogenic bacteria, where most cells are in contact with others of the same cell-type, would have a negative intermixing index II. Areas of high intermixing, where cells are mostly in contact with cells with differing cell-types, will have a positive intermixing index II.

### Statistical analyses

We performed all statistical analyses using R Studio Version 1.3.1073 (https://www.rstudio.com). We used parametric methods for all of our statistical tests and considered *P* < 0.05 to be statistically significant. We adjusted *P* for multiple comparison using the Benjamini-Hochberg method. We used the Shapiro-Wilk test to test for deviations from normality of our datasets. We considered *P* > 0.05 to validate the assumption of normality and found no evidence that our datasets significantly deviate from this assumption. We used the two-sample two-sided Welch test for all pair-wise comparisons, and we therefore did not make any assumptions regarding homogeneity of variances among our datasets. All sample sizes (n) reported in the results are the number of biological replicates.

## Data and code availability

All the source data and scripts required to reproduce the figures are publicly available on the Dryad Data repository at the following URL: https://datadryad.org/stash/share/SZITXkjfPBIgJKjEnFOfAWehPblsLbszTreUSE3IEsc. The CellModeller codes are also available publicly on Github CellModeller 4 repository, in the ‘Microbial Ecology Toolbox’ branch: https://github.com/HaseloffLab/CellModeller/tree/MicrobialEcologyToolbox/CellModeller. The code used to simulate the data is available in the ‘Examples/Ma2022/’ directory.

The scripts used to display the simulation output are in the ‘Scripts/’ directory. The code is additionally available at: https://github.com/antonkan/CellModeller4 in the ‘Microbial Ecology Toolbox’ branch.

## Acknowledgments

We thank Ella Flükiger for assistance with the experiments, Leo Eberl for the generous gift of plasmid pMA119, Malgorzata Kisielow and Anette Schütz from the Flow Cytometry Core Facility at ETH Zürich (https://facs.ethz.ch) for assistance with full-spectrum flow cytometry and data analysis, and Josep Ramoneda and Miaoxiao Wang for helpful discussions. Y.M. was supported by a grant from the Swiss National Science Foundation (31003A_176101) awarded to D.R.J.

## Author Contributions

Y.M. and D.R.J. conceived the research questions and designed the methodology and experiments. Y.M. performed the experiments. Y.M. and A.K. designed the computational model and performed simulations. Y.M. and D.R.J. analyzed and interpreted the data. D.R.J. initiated and coordinated the project. Y.M. and D.R.J. wrote the manuscript with input from A.K. All authors reviewed and approved the final version of the manuscript.

## APPENDIX

### Metabolic interactions determine the level of spatial intermixing between strains during microbial range expansion

We first performed range expansion experiments with consortia in the absence of plasmid pMA119 while imposing either a competitive (P+P) or a nitrite (NO_2_^-^) cross-feeding (P+C) interaction between the strains (Fig. S1A). We found that the nitrite cross-feeding interaction resulted in higher spatial intermixing than the competitive interaction (two-sample two-sided Welch test; P_A_ = 0.0026, n = 5), which is consistent with results reported in a previous study (Goldschmidt, Regoes et al. 2017). We next performed range expansion experiments with six consortia where one strain initially carried pMA119 (P*+P, P*+C, P+P* and P+C*) (Fig. S1B,C) or both strains initially carried pMA119 (P*+P* and P*+C*) (Fig. S1D) in the absence of antibiotic selection for pMA119. We found that the nitrite cross-feeding interaction consistently resulted in higher spatial intermixing than the competitive interaction (two-sample two-sided Welch test; P < 0.0007, n = 5) (Fig. S1A-D). We observed marginal loss of pMA119 (≈ 0) during the first week of expansion regardless of the type of metabolic interaction imposed (Fig. S2), demonstrating that pMA119 maintenance was strong under our experimental conditions and that pMA119 instability cannot confound our analyses. We therefore conclude that the type of metabolic interaction imposed between the strains does indeed determine the level of spatial intermixing that emerges during range expansion.

We next tested whether plasmid pMA119 itself affects the quantitative level of spatial intermixing that emerges between the strains during range expansion. To test this, we compared the levels of spatial intermixing across all the consortia for which we imposed the competitive interaction (P+P, P*+P, P+P* and P*+P*) (Fig. S1E) or the nitrite (NO_2_^-^) cross-feeding interaction (P+C, P*+C, P+C* and P*+C*) (Fig. S1F). We found that the levels of spatial intermixing are statistically identical regardless of whether the strains initially carried pMA119 for both the competitive interaction (one-way ANOVA test; P = 0.187, n = 5) (Fig. S1E) and the nitrite cross-feeding interaction (one-way ANOVA test; P = 0.096, n = 5) (Fig. S1F). Thus, pMA119 itself has no quantitative effect on the level of spatial intermixing that emerges during microbial range expansion.

**Appendix – Figure 1.**
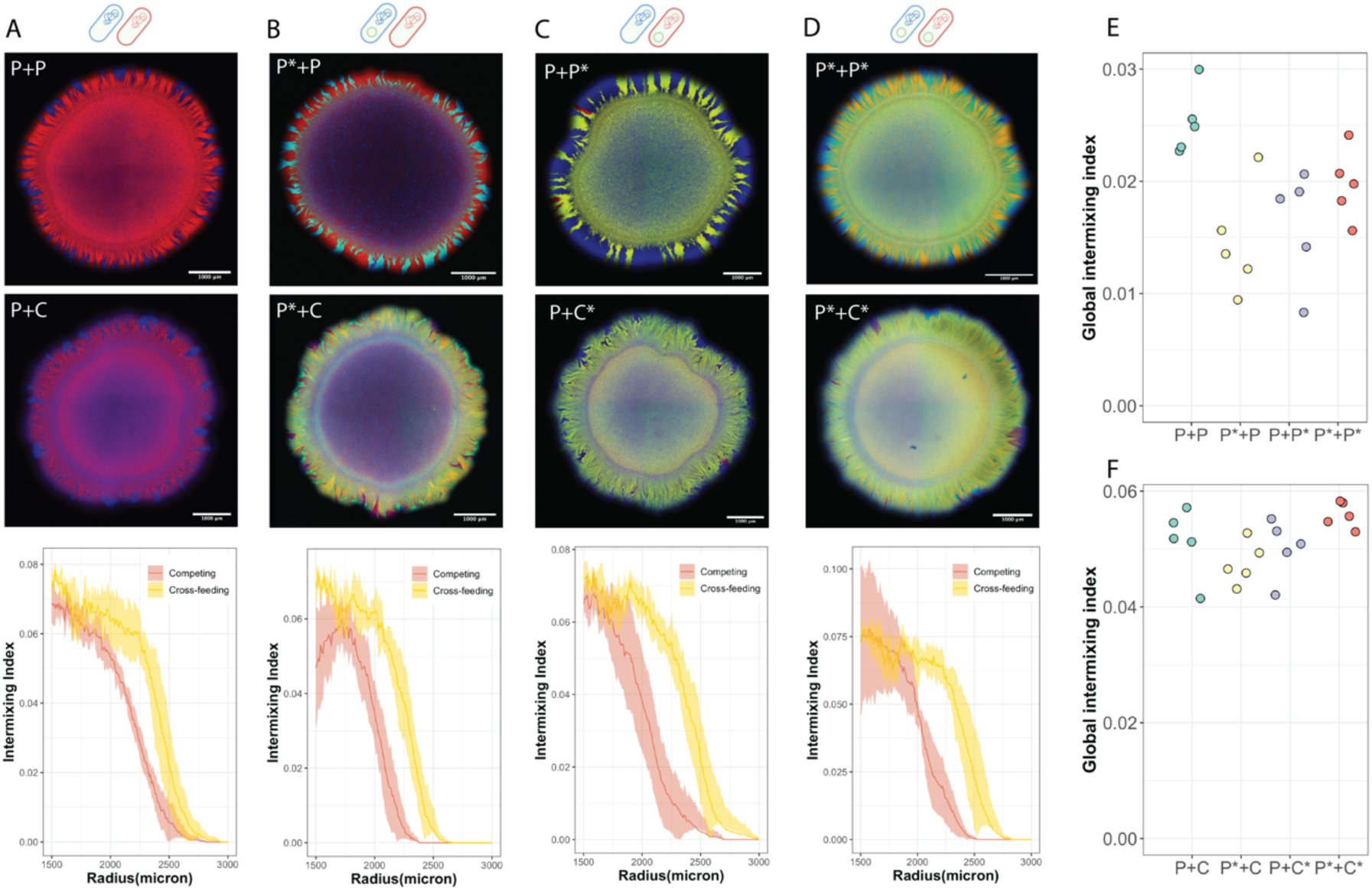
Metabolic interactions determine the level of spatial intermixing that emerges during range expansion. The two schematic cells above each image in **A-D** indicate the two strains used for each experiment. **A**, Neither strain initially carried plasmid pMA119 (P+P and P+C). **B**,**C**, One strain initially carried pMA119 (P*+P, P*+C, P+P*, and P+C*). **D**, Both strains initially carried pMA119 (P*+P* and P*+C*). For **A**-**D**, the top image is a representative range expansion for the competing interaction at the end of the experiment, the middle image is a representative range expansion for the nitrite (NO_2_^-^) cross-feeding interaction at the end of the experiment, and the bottom image is the intermixing index as a function of radius from the edge of the inoculation area (1500 µm) to the edge of the final expansion frontier (3000 µm) at radial increments of 10 µm. Data are presented for independent experimental replicates (n = 5) and the shaded regions are the standard deviations at each radial increment. **E,F**, Global intermixing index measured as the sum of intermixing indices across the expansion area at radial increments of 10 µm for consortia with **E** the competitive interaction or **F** the nitrite cross-feeding interaction.

**Appendix – Figure 2.**
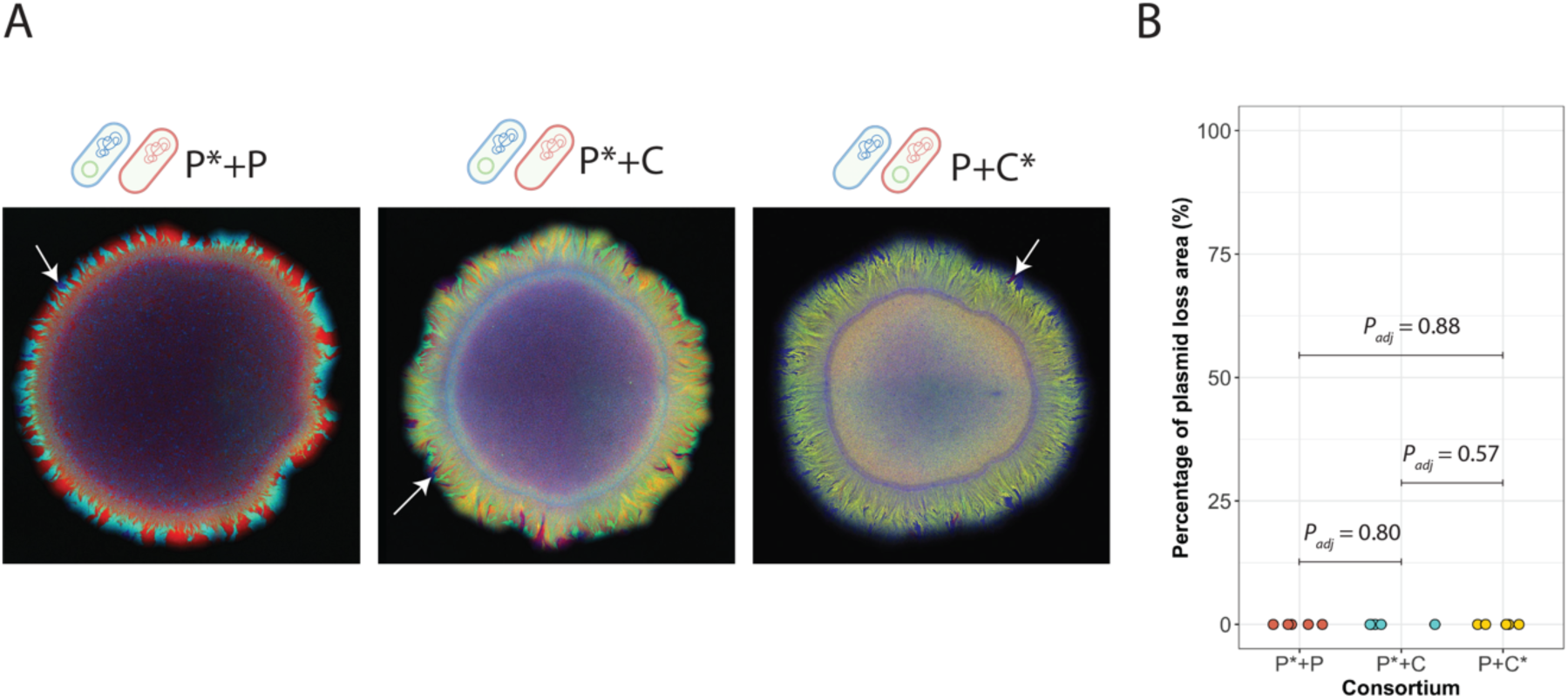
Quantification of plasmid loss after the first week of range expansion in the absence of antibiotic selection for pMA119. **A,** Representative images of range expansion after one week of range expansion using the same experimental design and procedures as for our main experiments. Consortium P*+P consisted of two producers that engaged in a competitive interaction, where one producer carried pMA119 (P*) while the other did not (P). Consortium P*+C consisted of a producer and consumer that engaged in a nitrite (NO_2_^-^) cross-feeding interaction, where the producer carried pMA119 (P*) while the consumer did not (C). Consortium P+C* also consisted of a producer and consumer that engaged in a nitrite (NO_2_^-^) cross-feeding interaction, but in this case the consumer carried pMA119 (C*) while the producer (P) did not. White arrows indicate regions where pMA119 was lost from the donor strain. These regions are identified as follows: if the producer loses pMA119 when it was the donor (P* to P), then blue regions emerge; If the consumer loses pMA119 when it was the donor (C* to C), then red regions emerge. **B,** Percentage of the producer or consumer that lost pMA119 when they were the donor. Each data point is the measurement for an independent experimental replicate (n = 5) and P_adj_ is the Benjamini-Hochberg-adjusted P for a two-sided two-sample Welch test.

**Appendix – Figure 3.**
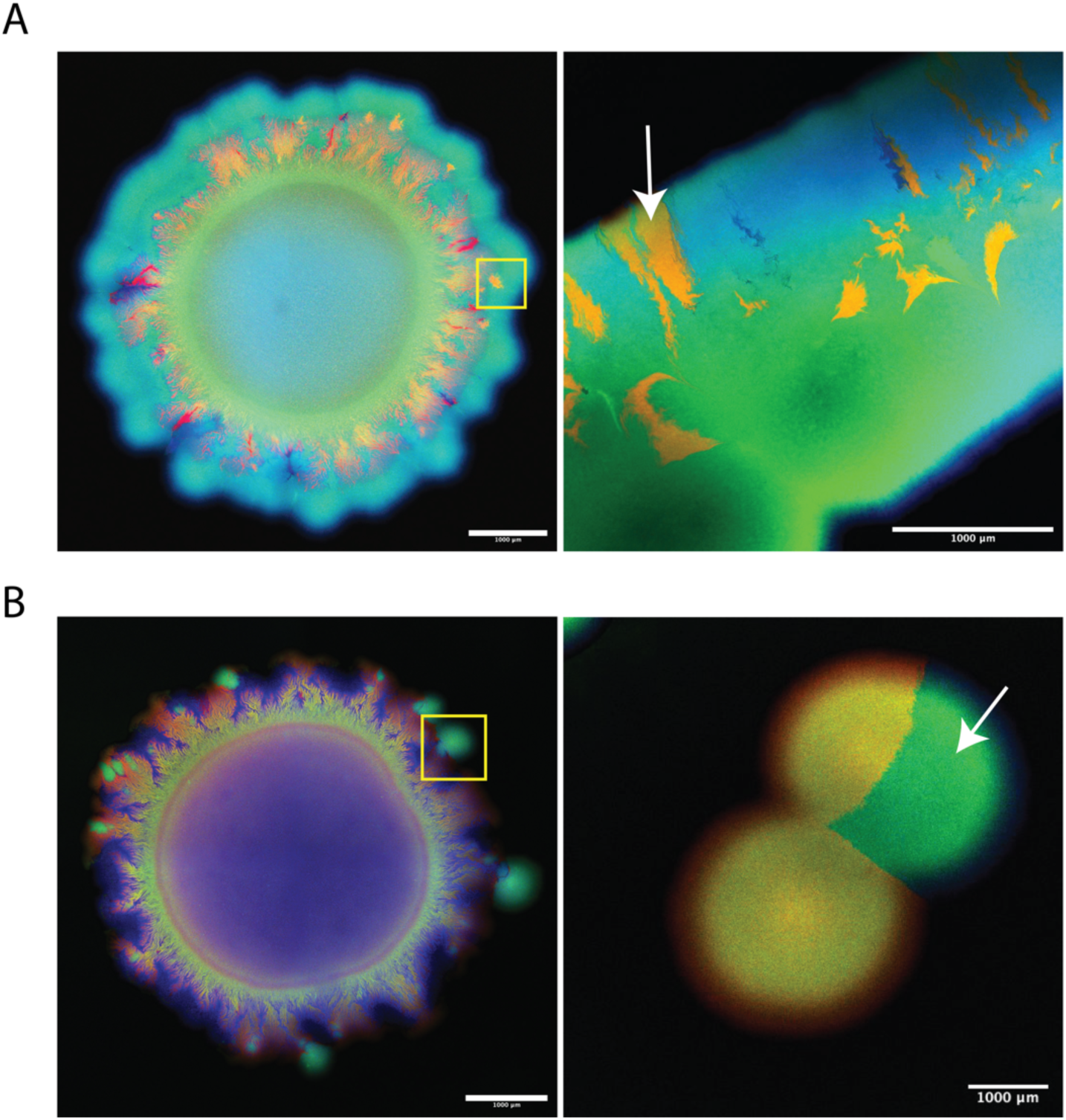
Verification of transconjugants after range expansion. Images are representative range expansions at the end of the experiment. **A**, The producer (pMA119 donor) expresses cyan and green fluorescent proteins while the consumer (potential pMA119 recipient) expresses red fluorescent protein. Transconjugants of the consumer appear yellow (indicated by the white arrow). **B**, The producer (potential pMA119 recipient) expresses cyan fluorescent protein while the consumer (pMA119 donor) expresses red and green fluorescent proteins. Transconjugants of the producer appear as green protrusions (indicated by the yellow frame). We sampled the transconjugant regions (green protrusions) with sterile toothpick and streaked the samples onto new LB agar plates containing kanamycin (50 µg ml^-1^) to verify that transconjugants

**Appendix – Figure 4.**
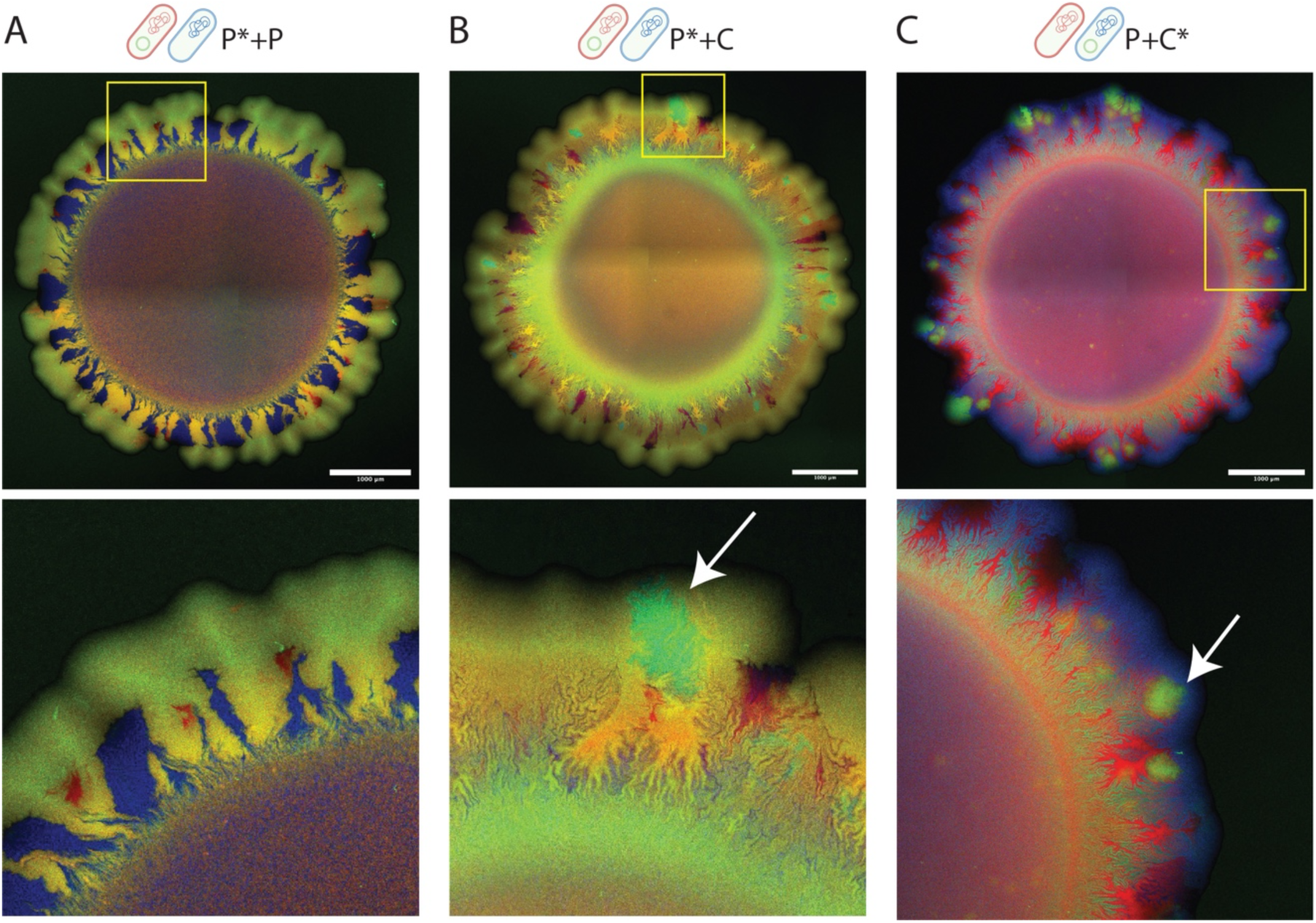
Color-swap experiments. We tested whether the chromosomally-located fluorescent protein-encoding gene expressed by the producer or consumer affects the emergence of transconjugants. **A-C**, The two schematic cells above each image indicate the two strains used for each experiment. Images are representative range expansions for consortia consisting of **A** two producers that engaged in a competitive interaction where one producer carried pMA119 while the other did not (P+P*), **B** a producer and consumer that engaged in a nitrite (NO_2_^-^) cross-feeding interaction where the producer carried pMA119 while the consumer did not (P*+C), and **C** a producer and consumer that engaged in a nitrite cross-feeding interaction where the consumer carried pMA119 while the producer did not (P+C*). White arrows indicate transconjugants, which we observed when we imposed the nitrite cross-feeding interaction but not the competitive interaction. Lower images are magnifications of the regions indicated by the yellow frames in the upper images. P, producer; C, consumer; *, pMA119 donor.

**Appendix – Figure 5.**
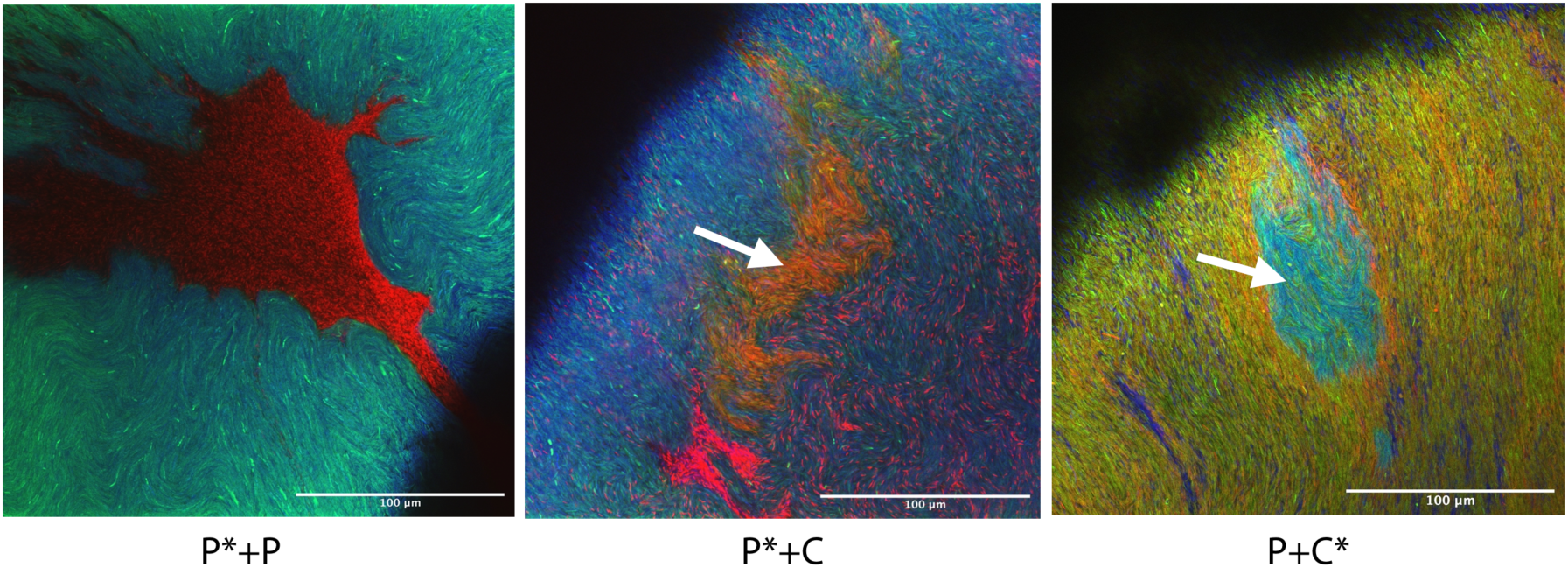
Different intermixing under 63-time magnification. Pictures were taken at the interfaces between two genotypes. In Group P* + P cyan strains (plasmid donors) and red strains (plasmid recipients) were completely segregated, generating clear borders; while cross-feeding group P*+C and Group P+C* show incredibly high mixing. White arrows represent transconjugant area under 63-time magnification.

**Appendix – Figure 6.**
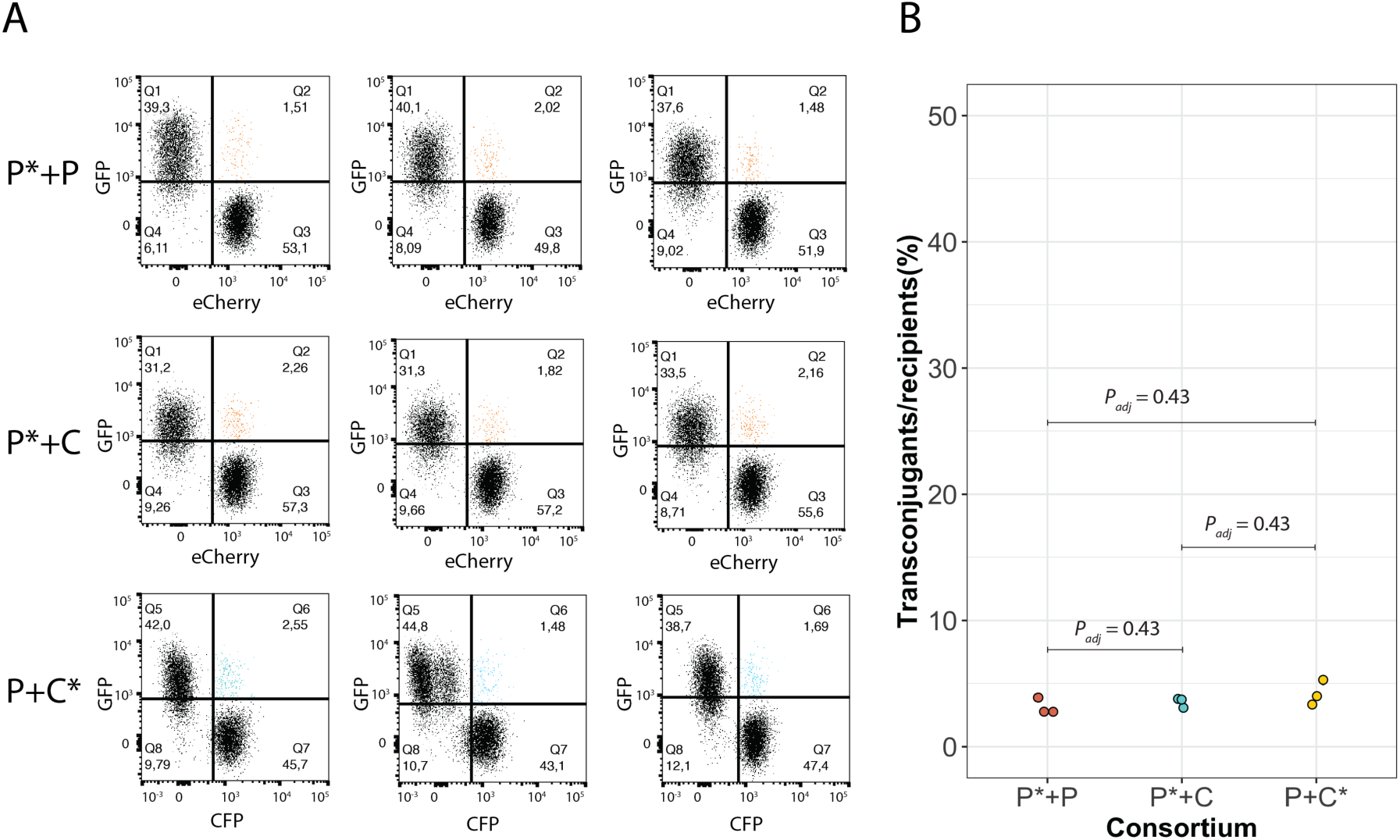
Quantification of transconjugants after filter mating under oxic conditions. We tested whether the producer and consumer have the same rate of plasmid pMA119 conjugation. **A**, Flow cytometry results for consortia consisting of two producers that engaged in a competitive interaction for oxygen where one producer carried pMA119 while the other did not (P*+P), a producer and consumer that engaged in a competitive interaction for oxygen where the producer carried pMA119 while the consumer did not (P*+C), and a producer and consumer that engaged in a competitive interaction for oxygen where the consumer carried pMA119 while the producer did not (P+C*). The upper left quadrants identify cells that only expressed green fluorescent protein, the bottom right quadrants identify cells that only expressed red fluorescent protein, and the upper right quadrants indicate transconjugants that expressed both fluorescent proteins. **B**, Quantification of the number of transconjugants per potential recipient for each consortium. Two-sample two-sided Welch tests showed no significant differences in the conjugation rates among the consortia. Each data point is the measurement for an independent experimental replicate (n = 3) and P_adj_ is the Benjamini-Hochberg-adjusted P for a two-sample two-sided Welch test. P, producer; C, consumer; *, pMA119 donor.

**Appendix – Figure 7.**
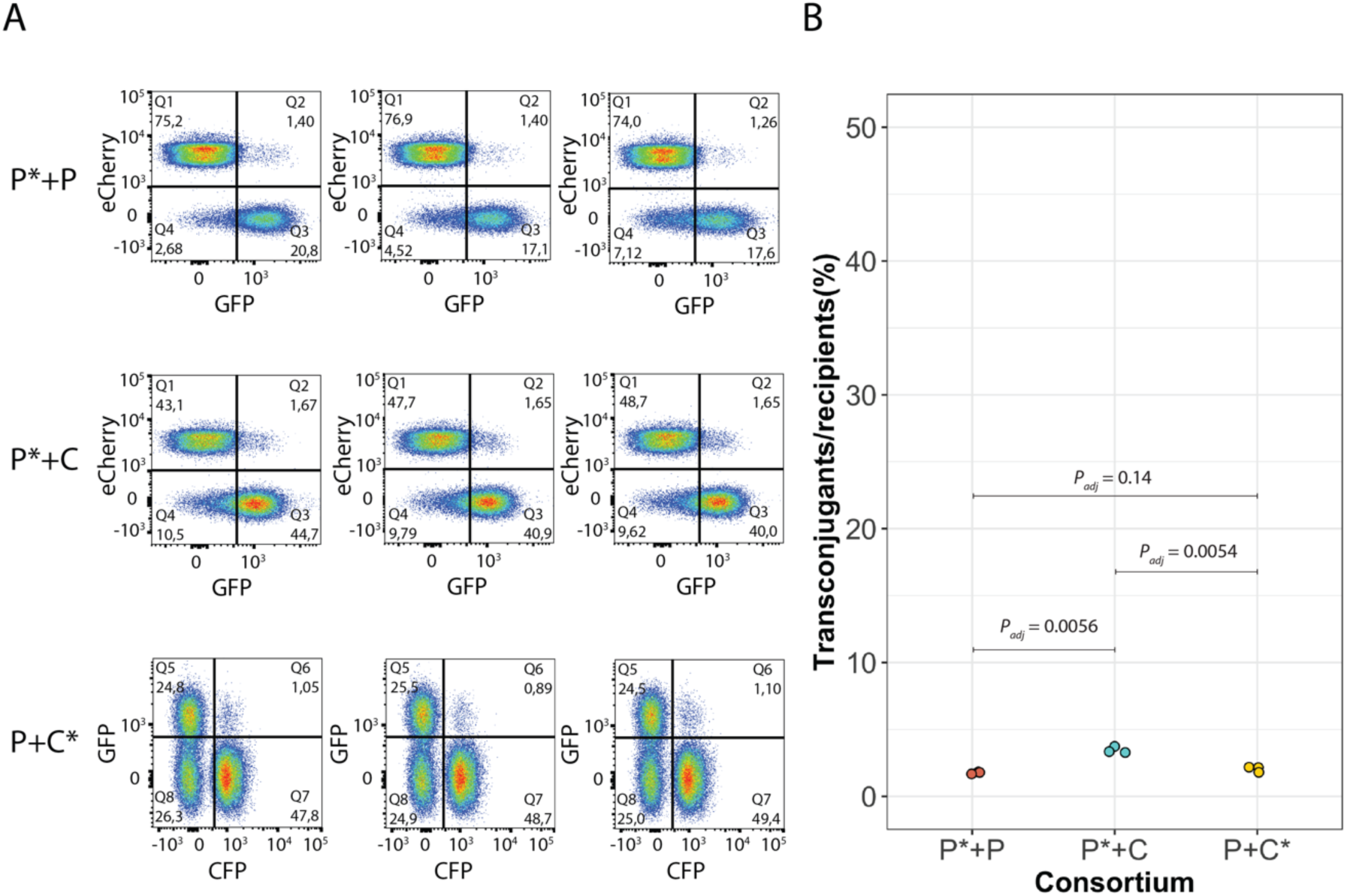
Quantification of transconjugants after growth in anoxic liquid medium. We tested whether the type of interaction imposed between the producer and consumer affects the extent of plasmid pMA119 conjugation. **A**, Flow cytometry results for consortia consisting of two producers that engaged in a competitive interaction where one producer carried pMA119 while the other did not (P+P*), a producer and consumer that engaged in a nitrite (NO_2_^-^) cross-feeding interaction where the producer carried pMA119 while the consumer did not (P*+C), and a producer and consumer that engage in a nitrite cross-feeding interaction where the consumer carried pMA119 while the producer did not (P+C*). The upper left quadrants identify cells that only expressed either red (for P*+P and P*+C) or green (for P+C*) fluorescent protein, the bottom right quadrants identify cells that only expressed either green (for P*+P and P*+C) or cyan (for P+C*) fluorescent protein, and the upper right quadrants identify transconjugants that expressed two fluorescent proteins. **B**, Quantification of the number of transconjugants per potential recipient for each consortium. Two-sample two-sided Welch tests showed significant differences but with small effect sizes. Each data point is the measurement for an independent experimental replicate (n = 3) and P_adj_ is the Benjamini-Hochberg-adjusted P for a two-sample two-sided Welch test. P, producer; C, consumer; *, pMA119 donor.

**Appendix – Figure 8.**
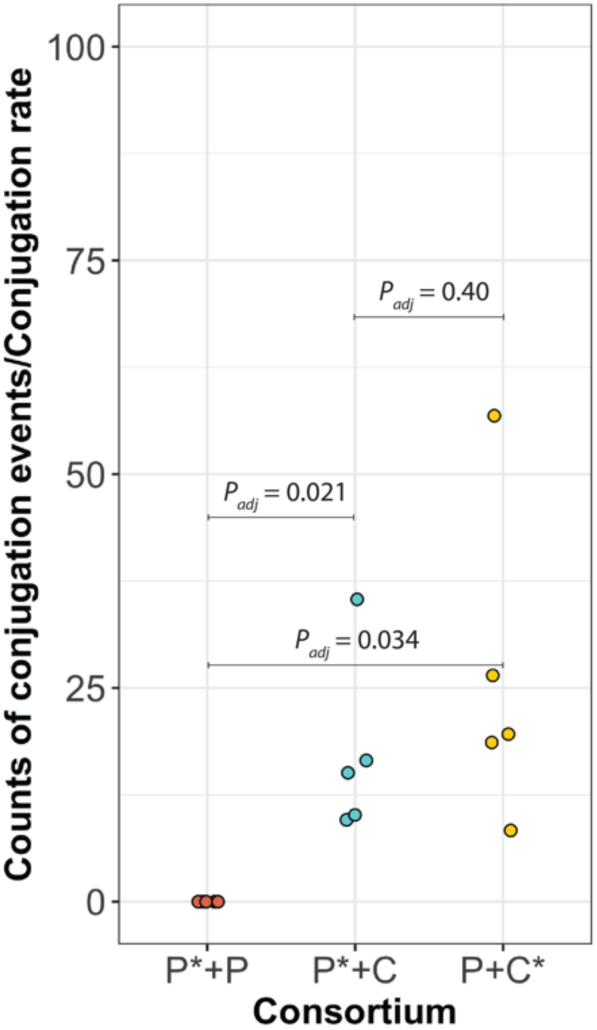
Counts of observed conjugation events normalized by the measured conjugation rate. The conjugation rates are those presented in Appendix – Figure 4. Data are for two producers that engaged in a competitive interaction where one producer carried plasmid pMA119 while the other did not (P+P*), a producer and consumer that engaged in a nitrite (NO_2_^-^) cross-feeding interaction where the producer carried pMA119 while the consumer did not (P*+C), and a producer and consumer that engaged in a nitrite cross-feeding interaction where the consumer carried pMA119 while the producer did not (P+C*). Each data point is the measurement for an independent experimental replicate (n = 5) and P_adj_ is the Benjamini-Hochberg-adjusted P for a two-sample two-sided Welch test. P, producer; C, consumer; *, pMA119 donor.

**Appendix – Figure 9.**
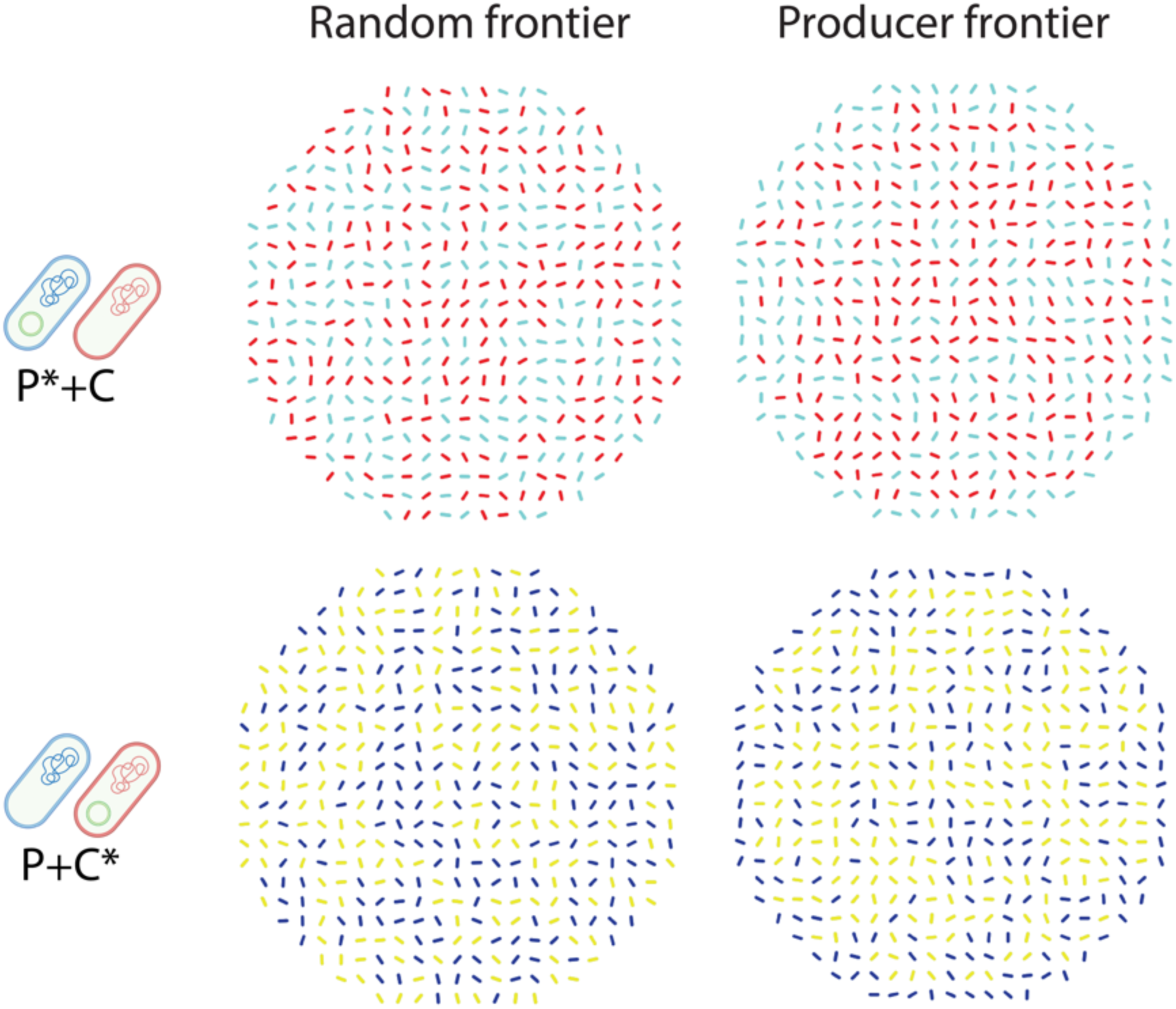
Initial spatial positioning of “Random frontier” and “Producer frontier”. The two schematic cells on the left side of the image indicate the two strains used for each experiment. Images are representative range expansions for consortia consisting of a producer and consumer that engaged in a nitrite (NO_2_^-^) cross-feeding interaction where the producer carried pMA119 while the consumer did not (P*+C), and a producer and consumer that engaged in a nitrite cross-feeding interaction where the consumer carried pMA119 while the producer did not (P+C*). As seen, in P*+C and P+C*, frontier is either dominated by both producer and consumer (Random frontier) or only by producer (Producer frontier).

**Appendix – Figure 10.**
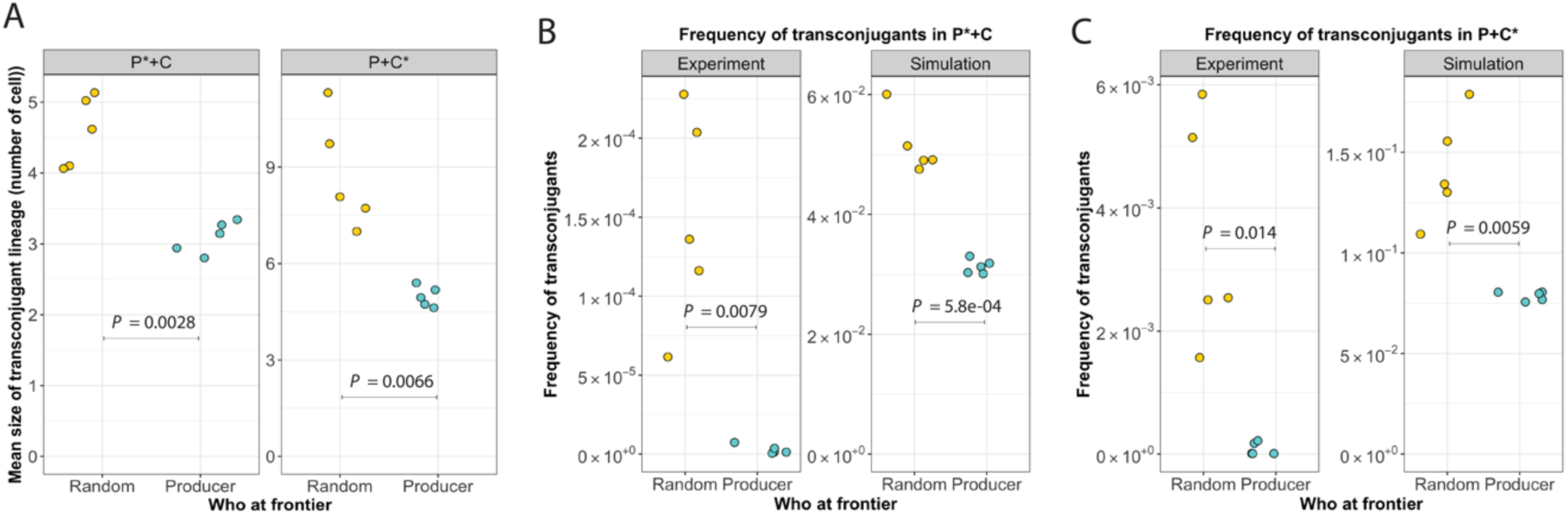
Effect of the spatial positionings on the proliferation of transconjugants during range expansion. **A,** Mean size of transconjugant lineage. **B,** Frequency of transconjugants in group P*+C. C, Frequency of transconjugants in group P+C*.

**Appendix – Figure 11.**
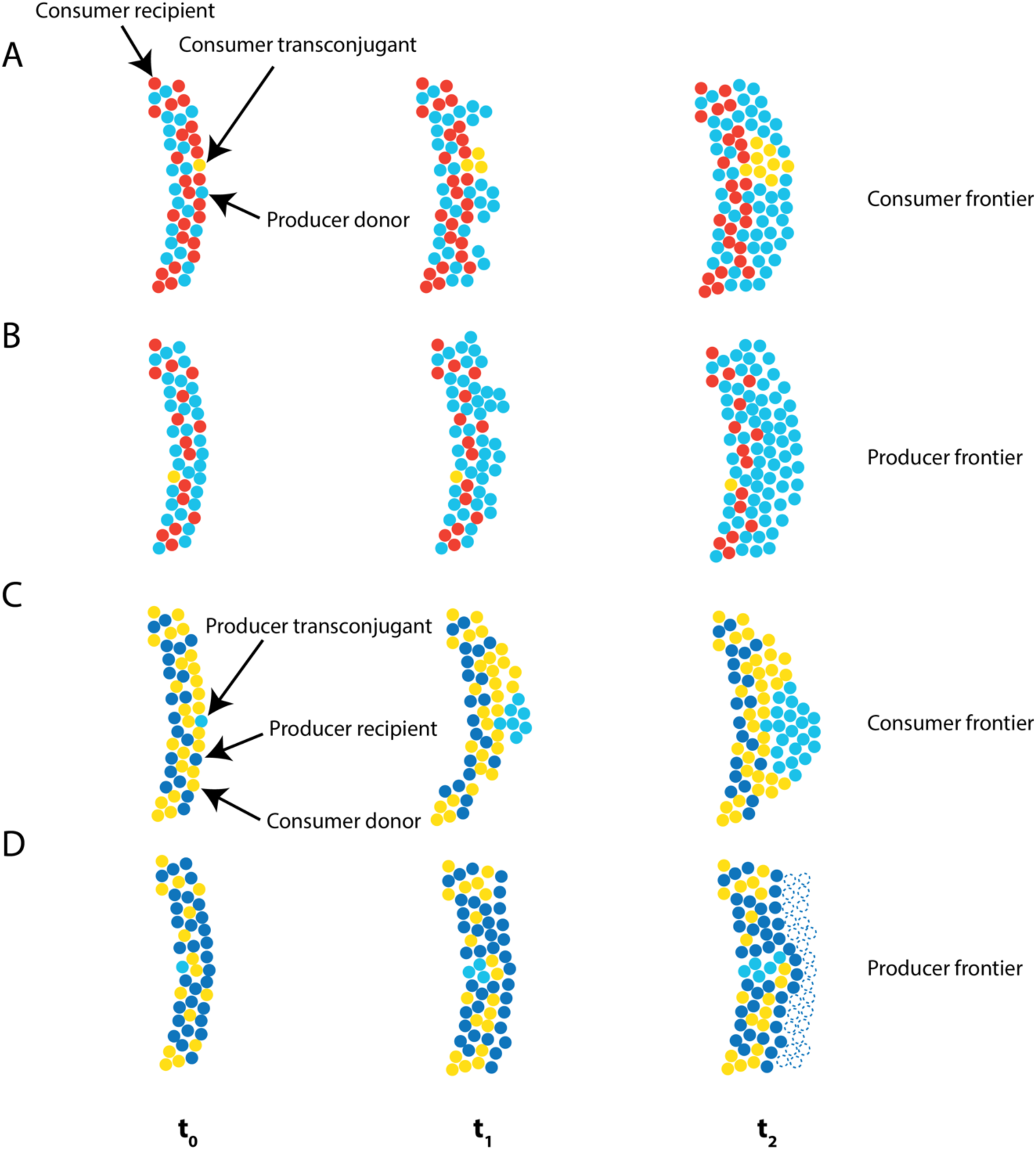
Schematic of how spatial positioning affects transconjugant proliferation. t_1_ and t_2_ indicate different expansion times, where t_0_ is early time, t_1_ is intermediate time, and t_2_ is late time. Depicted are only the frontiers of the expansion regions. Antibiotics were administered between t_0_ and t_1_. After antibiotic administration, only plasmid carriers could grow. **A**, The expansion frontier was primarily occupied by the consumer. The producer was the donor (cyan) while the consumer was the potential recipient (red). Transconjugants of the consumer are yellow. In this case, transconjugants of the consumer are more likely to emerge at the expansion frontier where resources are plentiful, and are thus more likely to proliferate. **B**, The expansion frontier was primarily occupied by the producer. The producer was the donor (cyan) while the consumer was the potential recipient (red). Transconjugants of the consumer are yellow. In this case, transconjugants of the consumer are less likely to emerge at the expansion frontier where resources are plentiful, and are thus less likely to proliferate. **C**, The expansion frontier was primarily occupied by the consumer. The producer was the potential recipient (blue) while the consumer was the donor (yellow). Transconjugants of the producer are cyan. In this case, transconjugants of the producer are more likely to emerge at the expansion frontier where resources are plentiful, and are thus more likely to proliferate. **D**, The expansion frontier was primarily occupied by the producer. The producer was the potential recipient (blue) while the consumer was the donor (yellow). Transconjugants of the producer are cyan. In this case, transconjugants of the producer are less likely to emerge at the expansion frontier where resources are plentiful, and are thus less likely to proliferate. Sensitive individuals of the producer will remain at the expansion frontier, but will not be able to grow in the presence of antibiotics (dashed circles).

**Appendix – Figure 12.**
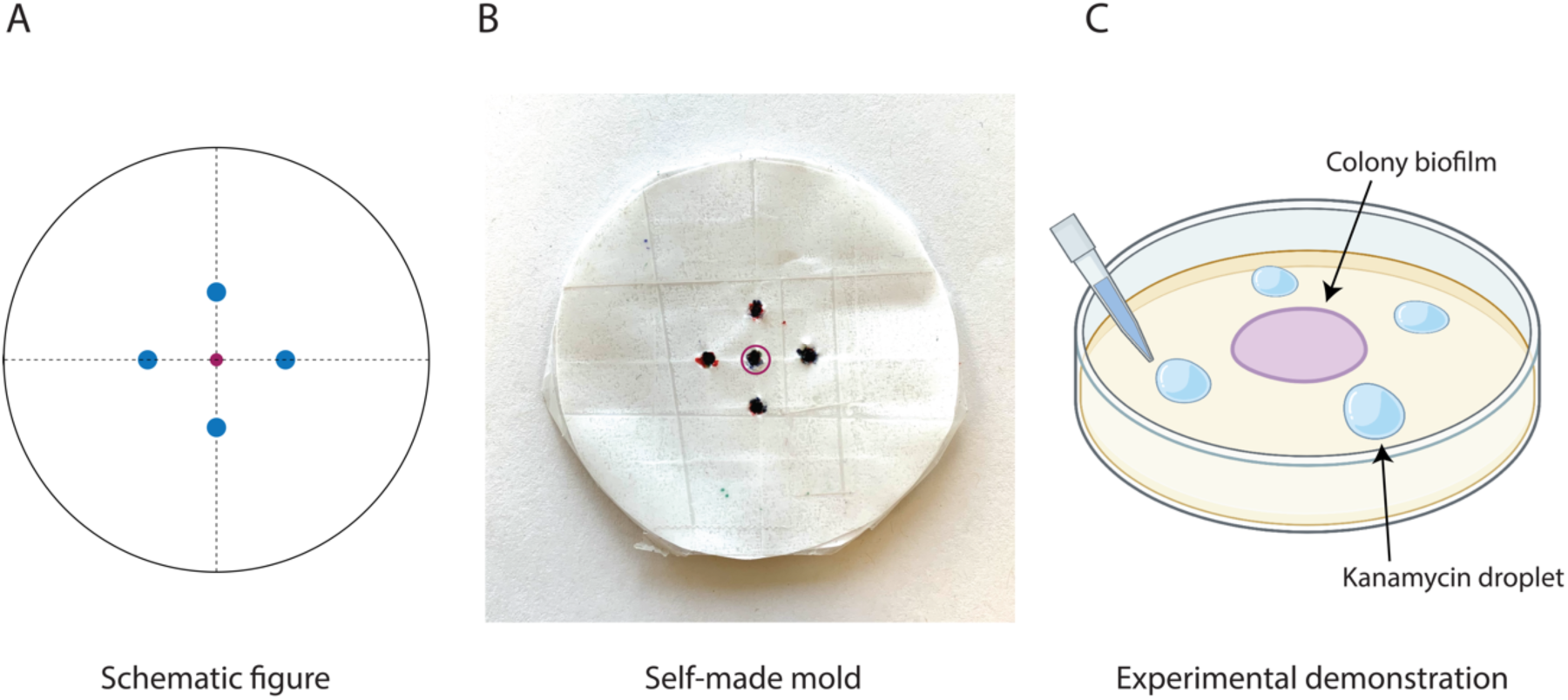
Mold used to apply kanamycin after one week of range expansion. **A**, Schematic design of the mold. The center point is the centroid of the range expansion. The four side points are equal distance from the centroid and are the points at which we added kanamycin. **B**, The mold shown that we used for this study. **C**, Schematic demonstration of the mold, where we placed the mold underneath the agar plate such that the center point marker aligns with the expansion centroid. We then deposited four droplets of kanamycin as point sources at each of the four side point markers. The final concentration of kanamycin within the agar plate was 50 µg/mL

**Appendix – Table 1.**
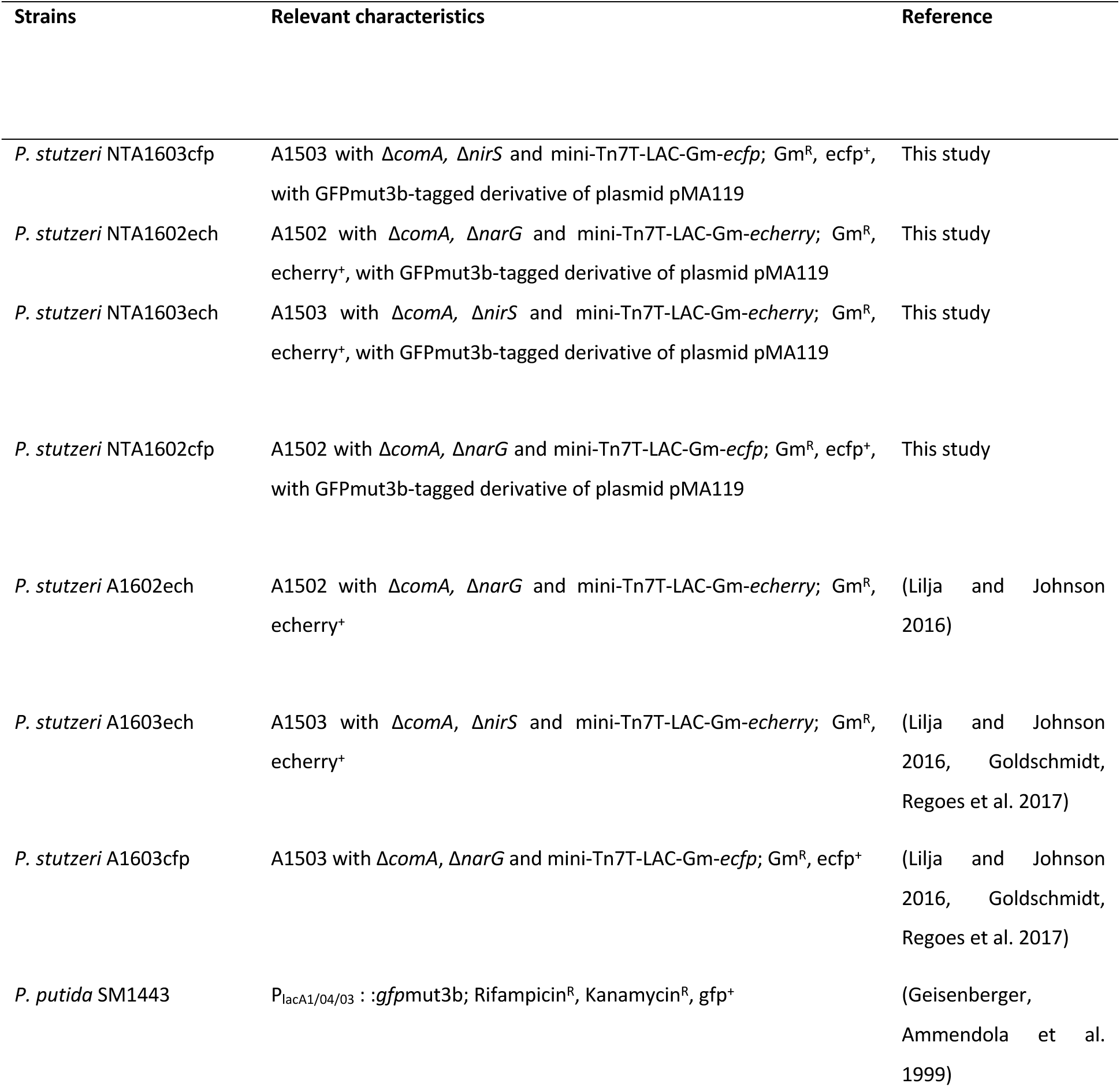
Specifications of the strains used in this study.

## References

Atis, S., B. T. Weinstein, A. W. Murray and D. R. Nelson (2019). “Microbial range expansions on liquid substrates.” Physical review X 9(2): 021058.

Babic, A., A. B. Lindner, M. Vulic, E. J. Stewart and M. Radman (2008). “Direct visualization of horizontal gene transfer.” Science 319(5869): 1533–1536.

Battin, T. J., K. Besemer, M. M. Bengtsson, A. M. Romani and A. I. Packmann (2016). “The ecology and biogeochemistry of stream biofilms.” Nature Reviews Microbiology 14(4): 251–263.

Ben-Jacob, E., I. Cohen and H. Levine (2000). “Cooperative self-organization of microorganisms.” Advances in Physics 49(4): 395–554.

Blanchard, A. E. and T. Lu (2015). “Bacterial social interactions drive the emergence of differential spatial colony structures.” BMC systems biology 9(1): 1–13.

Borer, B., D. Ciccarese, D. Johnson and D. Or (2020). “Spatial organization in microbial range expansion emerges from trophic dependencies and successful lineages.” Communications biology 3(1): 1–10.

Bosshard, L., I. Dupanloup, O. Tenaillon, R. Bruggmann, M. Ackermann, S. Peischl and L. Excoffier (2017). “Accumulation of Deleterious Mutations During Bacterial Range Expansions.” Genetics 207(2): 669–684.

Bosshard, L., S. Peischl, M. Ackermann and L. Excoffier (2019). “Mutational and Selective Processes Involved in Evolution during Bacterial Range Expansions.” Molecular Biology and Evolution 36(10): 2313–2327.

Bruellhoff, K., J. Fiedler, M. Möller, J. Groll and R. E. Brenner (2010). “Surface coating strategies to prevent biofilm formation on implant surfaces.” The International journal of artificial organs 33(9): 646–653.

Ciccarese, D., G. Micali, B. Borer, C. Ruan, D. Or and D. R. Johnson (2022). “Rare and localized events stabilize microbial community composition and patterns of spatial self-organization in a fluctuating environment.” The ISME Journal: 1–11.

Ciccarese, D., A. Zuidema, V. Merlo and D. R. Johnson (2020). “Interaction-dependent effects of surface structure on microbial spatial self-organization.“ Philosophical Transactions of the Royal Society B 375(1798): 20190246.

Excoffier, L., M. Foll and R. J. Petit (2009). “Genetic consequences of range expansions.” Annual Review of Ecology, Evolution, and Systematics 40: 481–501.

Fei, C., S. Mao, J. Yan, R. Alert, H. A. Stone, B. L. Bassler, N. S. Wingreen and A. Košmrlj (2020). “Nonuniform growth and surface friction determine bacterial biofilm morphology on soft substrates.” Proceedings of the National Academy of Sciences 117(14): 7622–7632.

Ferreira, T. A., A. V. Blackman, J. Oyrer, S. Jayabal, A. J. Chung, A. J. Watt, P. J. Sjöström and D. J. Van Meyel (2014). “Neuronal morphometry directly from bitmap images.” Nature methods 11(10): 982–984.

Flemming, H.-C. and J. Wingender (2010). “The biofilm matrix.” Nature reviews microbiology 8(9): 623–633.

Geisenberger, O., A. Ammendola, B. B. Christensen, S. Molin, K.-H. Schleifer and L. Eberl (1999). “Monitoring the conjugal transfer of plasmid RP4 in activated sludge and in situ identification of the transconjugants.” FEMS microbiology letters 174(1): 9–17.

Giometto, A., D. R. Nelson and A. W. Murray (2018). “Physical interactions reduce the power of natural selection in growing yeast colonies.” Proceedings of the National Academy of Sciences 115(45): 11448–11453.

Goldschmidt, F., L. Caduff and D. R. Johnson (2021). “Causes and consequences of pattern diversification in a spatially self-organizing microbial community.” The ISME Journal: 1–12.

Goldschmidt, F., R. R. Regoes and D. R. Johnson (2017). “Successive range expansion promotes diversity and accelerates evolution in spatially structured microbial populations.” The ISME journal 11(9): 2112–2123.

Gralka, M. and O. Hallatschek (2019). “Environmental heterogeneity can tip the population genetics of range expansions.” Elife 8.

Gralka, M., F. Stiewe, F. Farrell, W. Möbius, B. Waclaw and O. Hallatschek (2016). “Allele surfing promotes microbial adaptation from standing variation.” Ecology letters 19(8): 889–898.

Hall-Stoodley, L., J. W. Costerton and P. Stoodley (2004). “Bacterial biofilms: from the natural environment to infectious diseases.” Nature reviews microbiology 2(2): 95–108.

Hallatschek, O., P. Hersen, S. Ramanathan and D. R. Nelson (2007). “Genetic drift at expanding frontiers promotes gene segregation.” Proceedings of the National Academy of Sciences 104(50): 19926–19930.

Hallatschek, O. and D. R. Nelson (2010). “Life at the front of an expanding population.” Evolution 64(1): 193–206.

Hayes, C. S., S. K. Aoki and D. A. Low (2010). “Bacterial contact-dependent delivery systems.” Annu Rev Genet 44: 71–90.

Kan, A., I. Del Valle, T. Rudge, F. Federici and J. Haseloff (2018). “Intercellular adhesion promotes clonal mixing in growing bacterial populations.” Journal of The Royal Society Interface 15(146): 20180406.

Kayser, J., C. F. Schreck, Q. Yu, M. Gralka and O. Hallatschek (2018). “Emergence of evolutionary driving forces in pattern-forming microbial populations.” Philosophical Transactions of the Royal Society B: Biological Sciences 373(1747): 20170106.

Lilja, E. E. and D. R. Johnson (2016). “Segregating metabolic processes into different microbial cells accelerates the consumption of inhibitory substrates.” The ISME journal 10(7): 1568–1578.

Lilja, E. E. and D. R. Johnson (2017). “Metabolite toxicity determines the pace of molecular evolution within microbial populations.” BMC evolutionary biology 17(1): 1–12.

Mitri, S., E. Clarke and K. R. Foster (2016). “Resource limitation drives spatial organization in microbial groups.” The ISME journal 10(6): 1471–1482.

Momeni, B., K. A. Brileya, M. W. Fields and W. Shou (2013). “Strong inter-population cooperation leads to partner intermixing in microbial communities.” elife 2: e00230.

Momeni, B., A. J. Waite and W. Shou (2013). “Spatial self-organization favors heterotypic cooperation over cheating.” Elife 2: e00960.

Müller, M. J., B. I. Neugeboren, D. R. Nelson and A. W. Murray (2014). “Genetic drift opposes mutualism during spatial population expansion.” Proceedings of the National Academy of Sciences 111(3): 1037–1042.

Nadell, C. D., K. Drescher and K. R. Foster (2016). “Spatial structure, cooperation and competition in biofilms.” Nature Reviews Microbiology 14(9): 589–600.

Nadell, C. D., K. R. Foster and J. B. Xavier (2010). “Emergence of spatial structure in cell groups and the evolution of cooperation.” PLoS computational biology 6(3): e1000716.

Nobile, C. J. and A. D. Johnson (2015). “Candida albicans biofilms and human disease.” Annual review of microbiology 69: 71–92.

Rodríguez Amor, D. and M. Dal Bello (2019). “Bottom-up approaches to synthetic cooperation in microbial communities.” Life 9(1): 22.

Ruan, C., J. Ramoneda, G. Chen, D. R. Johnson and G. Wang (2021). “Evaporation-induced hydrodynamics promote conjugation-mediated plasmid transfer in microbial populations.” ISME Communications 1(1): 54.

Rudge, T. J., F. Federici, P. J. Steiner, A. Kan and J. Haseloff (2013). “Cell Polarity-Driven Instability Generates Self-Organized, Fractal Patterning of Cell Layers.” ACS Synthetic Biology 2(12): 705–714.

Rudge, T. J., P. J. Steiner, A. Phillips and J. Haseloff (2012). “Computational modeling of synthetic microbial biofilms.” ACS synthetic biology 1(8): 345–352.

Sharma, A. and K. B. Wood (2021). “Spatial segregation and cooperation in radially expanding microbial colonies under antibiotic stress.” The ISME Journal: 1–15.

Sijbesma, W. F., J. S. Almeida, M. A. Reis and H. Santos (1996). “Uncoupling effect of nitrite during denitrification by Pseudomonas fluorescens: An in vivo 31P-NMR study.” Biotechnology and bioengineering 52(1): 176–182.

Singh, R., D. Paul and R. K. Jain (2006). “Biofilms: implications in bioremediation.” Trends in Microbiology 14(9): 389–397.

Smith, W. P., Y. Davit, J. M. Osborne, W. Kim, K. R. Foster and J. M. Pitt-Francis (2017). “Cell morphology drives spatial patterning in microbial communities.” Proceedings of the National Academy of Sciences 114(3): E280–E286.

Sørensen, S. J., M. Bailey, L. H. Hansen, N. Kroer and S. Wuertz (2005). “Studying plasmid horizontal transfer in situ: a critical review.” Nat Rev Microbiol 3(9): 700–710.

Stalder, T. and E. Top (2016). “Plasmid transfer in biofilms: a perspective on limitations and opportunities.” NPJ Biofilms Microbiomes 2: 16022–.

Stalder, T. and E. Top (2016). Plasmid transfer in biofilms: a perspective on limitations and opportunities. NPJ Biofilms Microbiomes 2: 16022.

Taheri-Araghi, S., S. Bradde, J. T. Sauls, N. S. Hill, P. A. Levin, J. Paulsson, M. Vergassola and S. Jun (2015). “Cell-size control and homeostasis in bacteria.” Current biology 25(3): 385–391.

Tecon, R., A. Ebrahimi, H. Kleyer, S. Erev Levi and D. Or (2018). “Cell-to-cell bacterial interactions promoted by drier conditions on soil surfaces.” Proceedings of the National Academy of Sciences 115(39): 9791–9796.

Tecon, R. and D. Or (2017). “Cooperation in carbon source degradation shapes spatial self-organization of microbial consortia on hydrated surfaces.” Scientific reports 7(1): 1–11.

Thomas, C. M. and K. M. Nielsen (2005). “Mechanisms of, and barriers to, horizontal gene transfer between bacteria.” Nat Rev Microbiol 3(9): 711–721.

Tolker-Nielsen, T. and S. Molin (2000). “Spatial organization of microbial biofilm communities.” Microbial ecology 40(2): 75–84.

Weinstein, B. T., M. O. Lavrentovich, W. Möbius, A. W. Murray and D. R. Nelson (2017). “Genetic drift and selection in many-allele range expansions.” PLoS computational biology 13(12): e1005866.

Xiong, L., Y. Cao, R. Cooper, W.-J. Rappel, J. Hasty and L. Tsimring (2020). “Flower-like patterns in multi-species bacterial colonies.” Elife 9: e48885.

Yanni, D., P. Márquez-Zacarías, P. J. Yunker and W. C. Ratcliff (2019). “Drivers of spatial structure in social microbial communities.” Current biology 29(11): R545–R550.

Yu, Q., M. Gralka, M. C. Duvernoy, M. Sousa, A. Harpak and O. Hallatschek (2021). “Mutability of demographic noise in microbial range expansions.” Isme j 15(9): 2643–2654.

Zhou, Y., A. Oehmen, M. Lim, V. Vadivelu and W. J. Ng (2011). “The role of nitrite and free nitrous acid (FNA) in wastewater treatment plants.” Water research 45(15): 4672–4682.

